# Modulatory effect of non-invasive prefrontal brain stimulation on risk-taking behaviours in humans: preliminary insight on the influence of personality traits

**DOI:** 10.1101/2022.11.22.517525

**Authors:** Ona Martin de la Torre, Antoni Valero-Cabré, David Gallardo-Pujol, Diego Redolar-Ripoll

## Abstract

We investigated the modulatory effects of cathodal High-Definition transcranial Direct Current Stimulation (HD-tDCS) on the left dorsolateral prefrontal cortex (DLPFC) and the left ventrolateral prefrontal cortex (VLPFC) on risk-taking.

**Methods:** Thirty-four healthy adults underwent 3 independent cathodal HD-tDCS interventions (DLPFC, VLPFC, sham) delivered in counterbalanced order during the performance of the balloon analogue risk task (autoBART). Participants were clustered post-hoc in 3 separate personality profiles according to the HEXACO-60 and the Dark Triad dirty dozen and we reanalysed the data.

**Results:** Dorsal prefrontal cathodal HD-tDCS significantly modulated autoBART performance rendering participants less prone to risk-taking (i.e., more conservative) under left DLPFC HD-tDCS compared to left VLPFC or sham stimulation. The re-analysis of the same dataset, taking into consideration personality traits, suggested specific effects in impulsive-disinhibited and normative participants for DLPFC and VLPFC stimulation, respectively. Specifically, we saw that participants classified as impulsive-disinhibited were more affected by HD-tDCS left DLPFC stimulation than other profiles.

**Conclusions:** Both, dorsal and ventral prefrontal active HD-tDCS decrease risk-taking behaviour compared to sham stimulation. Importantly, such effects are likely influenced by personality traits (impulsive disinhibited vs normative) exhibited by the participants.

**Highlights:**

- We investigated the effect of dorsal and ventral prefrontal HD-tDCS on risk-taking.
- We considered post-hoc, the influence of individual personality differences.
- Dorsal and ventral Prefrontal cathodal HD-tDCS decreased risk-taking behaviour.
- Left cathodal DLPFC HD-tDCS decreased risk propensity in impulsive-disinhibited participants.
- Left cathodal VLPFC HD-tDCS decreased risk propensity in normative personality participants.

## 1. INTRODUCTION

As part of our daily life, we are prompt to constantly make decisions, involving important and complex choices like planning our professional career, or trivial ones such as choosing tea or coffee for breakfast. Some everyday decisions are rather simple and require little cognitive effort, whereas complex ones require detailed analysis and profound value judgements to select among options leading to the desired outcomes (Busemeyer *et al*., 2019). Cognitive control functions subtended by prefrontal systems allow us to adequately integrate our own perceptions, goals, and motivations and weigh them with prior knowledge and current information to ultimately take adaptive value-based decisions (Obeso *et al*., 2013).

Cognitive neuroscience has largely explored the neural correlates involved in value-based decision-making in healthy individuals (Poudel *et al*., 2017) or in brain damage (Bechara *et al*., 1994; Bechara *et al*., 2000) and psychiatric conditions (Ernst *et al*., 2003; Glenn *et al*., 2009). This processes are instantiated by dedicated brain systems (Ernst and Paulus, 2005; Gold and Shadlen, 2007; Peter N.C. Mohr *et al*., 2010; Khani and Rainer, 2016; Lee and Seo, 2016; Kurikawa *et al*., 2018; Atiya *et al*., 2019) subtending a variety of sub-processes including the evaluation of available options (Gutnik *et al*., 2006; Bossaerts and Murawski, 2015; Szrek, 2017), the ponderation of gains and losses (Tom *et al*., 2007; Zhang *et al*., 2017; Sokol-Hessner and Rutledge, 2019), the computation of outcome probabilities (Troffaes, 2007; Huang *et al*., 2011; Chen *et al*., 2012), and the consideration of uncertainty and decision confidence (Pushkarskaya *et al*., 2015; Kurikawa *et al*., 2018; Atiya *et al*., 2019).

A value-based decision-making type, quite common in real-life decisions, is associated with risk-taking (Megías *et al*., 2018). Multiple regions of the prefrontal cortex such as dorsolateral areas (Ernst *et al*., 2002; F *et al*., 2002; Brand *et al*., 2005; Fellows and Farah, 2005; Labudda *et al*., 2008; Steinberg, 2008), ventrolateral regions (Fellows and Farah, 2007; Hampshire *et al*., 2008; Baxter *et al*., 2009; Guo *et al*., 2013; Domenech and Koechlin, 2015; Chung *et al*., 2017; Wearne, 2018), areas of the orbitofrontal cortex (R D Rogers *et al*., 1999; R. D. Rogers *et al*., 1999; Manes *et al*., 2002; Clark *et al*., 2003; Ernst *et al*., 2004; Fukui *et al*., 2005), parietal areas (Coutlee *et al*., 2016), regions of the anterior cingulate (Ernst *et al*., 2002; Labudda *et al*., 2008; Lawrence *et al*., 2009), and insular cortices (Ernst *et al*., 2002; Clark *et al*., 2008; Smith *et al*., 2009; Bar-On *et al*., 2013) have been involved in cost benefit evaluations of potential risks and rewards (St. Onge and Floresco, 2010).

In agreement with such anatomical basis, Parkinson (Mimura *et al*., 2006), schizophrenia (Sterzer *et al*., 2019), frontal damage patients (Bechara *et al*., 1994; Bechara *et al*., 1999), chronic drug-users (Clark and Robbins, 2002; Ekhtiari *et al*., 2017) and also pathological gamblers (Bechara *et al*., 1994; de Ruiter *et al*., 2009) with alterations of prefrontal dopamine systems show perturbations of risk-based decision-making. Nonetheless, the specific causal role and functional contributions of dorsal and ventral prefrontal cortical regions to such processes remain to be further explored.

Transcranial magnetic stimulation (TMS) and transcranial direct current stimulation (tDCS) allow for the direct manipulation of activity patterns in cortical regions. Both techniques have been used to causally dissect the causal contribution of prefrontal cortex sub-regions to different subprocesses involved in risk decision-making (Pascual-Leone *et al*., 1999; Nevler and Ash, 2015).

TMS evidence suggested that disruption of either the left lateral prefrontal cortex (LPFC) (Figner *et al*., 2010) or the right dorsolateral prefrontal cortex (DLPFC) (Knoch *et al*., 2006; Tulviste and Bachmann, 2019) increase impulsivity and risk-taking behaviour. Additionally, right anodal/left cathodal dual tDCS on the DLPFC increases response confidence (Minati *et al*., 2012), reduces risk-taking behaviour (Fecteau, Knoch, *et al*., 2007; Cheng and Lee, 2016), and unpromoted risk-taking strategies that avoid the risk of no reward (Ota *et al*., 2019). Moreover, the opposite montage, anodal left DLPFC combined with cathodal right DLPFC stimulation has shown to increase, in elderly individuals, the likelihood of risk-taking options (Paulo S. Boggio *et al*., 2010). Lastly, both, right anodal/left cathodal and left anodal/right cathodal tDCS montages on the DLPFC have led to less risk-taking behaviour (Fecteau, Pascual-Leone, *et al*., 2007). Nonetheless, in contrast, other authors have shown that the same montages, compared to sham stimulation, increased risk-taking (Ye, Chen, Huang, Wang, Jia, *et al*., 2015), with an asymmetric effect for the right/left DLPFC when participants confronted losses and gains.

Regarding the ventrolateral prefrontal cortex (VLPFC), different neuroimaging studies have shown the implication of this region in uncertain decision-making (Fellows and Farah, 2007), and reported a VLPFC activation decreases associated with a longitudinal decline in self-reported risk behaviour during adolescence (Qu *et al*., 2015). Other research has shown reduced risk-taking behaviour when mothers were present (Telzer *et al*., 2015) and additionally, strong activation of this same region for conditions of high risk. Furthermore, fMRI studies have linked the modulation of the VLPFC to negative feedback processing in adults and children (van Leijenhorst *et al*., 2006) and to negative emotions (Vergallito *et al*., 2018). Taken together, such a set of diverse correlational evidence suggests the role of VLPFC in regulating negative emotions that could affect risk decision-making. Nonetheless, to date no study using non-invasive brain stimulation (NIBS) has added causality to some above-drawn associations.

On the basis of the above-mentioned literature, here we examined the modulatory effects of HD-tDCS on risk-taking. We hypothesized that cathodal HD-transcranial direct current stimulation (HD-tDCS) of the left DLPFC and the left VLPFC would decrease risk propensity. In a second step, given accounts that personality influences human decision-making (Neisser, 1967; Jalajas and Pullaro, 2018), reports that psychopathy scores correlate with the percentage of risky decisions made during the Cambridge decision-making task (Sutherland and Fishbein, 2017), as well as the fact that life outcomes can be linked to personality traits (Soto, 2019), personality profiles were taken into consideration in a reanalysis of the same dataset.

## 2. MATERIALS AND METHODS

### 2.1. Participants

A total of 34 right-handed healthy college-educated volunteers participated in the study (21 females and 13 males, mean age 29.21±9.72 years). None were taking medication of any kind, showed previous history of neurological disorders, or psychiatric illness, neither of drug or alcohol abuse. Additionally, all participants met internationally-established safety criteria to receive tDCS (Nitsche *et al*., 2003; Bikson *et al*., 2017). The local ethics committees of the Open University of Catalonia (UOC) and the institutional review board (IRB 00003099) of the University of Barcelona approved the study, complying with the principles of the Declaration of Helsinki. All participants provided written informed consent and received an established financial compensation for their participation at the end of the study.

### 2.2. Experimental design and general procedure

Using a cross over-counterbalanced design, we aimed at assessing the online impact of a single session of tDCS delivered to the DLPFC or the VLPFC and compared to a sham tDCS condition, on risk taking behaviour. Accordingly, the study consisted of three HD-tDCS sessions delivered across 3 independent sessions, with DLPFC, VLPFC and a sham condition (see below for full details on electrode localisation) applied in a randomized and counterbalanced order. Following prior evidence showing that the effects of a single stimulation session did not endure beyond 1 hour, sessions were sufficiently spaced to avoid unlikely carry-over effects (Nitsche *et al*., 2008).

To explore whether the HD-tDCS-induced effect on risk decision-making depended on stimulation intensity, participants were randomly assigned to one of two different intensity levels (1.5mA or 2mA) maintained during the whole session. Previous studies showed that 2 mA tDCS does not necessarily yield larger effects than 1.5 mA tDCS in healthy participants (Ho *et al*., 2016; Jamil *et al*., 2017). Jamil and colleagues investigated tDCS intensity ranges from 0.5 mA to 2.0 mA for anodal and cathodal tDCS and lower intensities of tDCS shows equal or greater results in motor-cortical excitability. Therefore, we decided to treat intensity as a second independent variable, since there has not been any research that dealt with different intensities on a risk decision-making task. And to ascertain if the modulatory effects could be modulated in magnitude, ranging from moderate/mild effects to strong/intense effects.

During each session, participants performed the Balloon Analogue Risk Task (BART), a computerized risk decision-making task. We made sure prior to the experimental sessions that the participants would be able to complete the task under 20 minutes (the duration of the HD-tDCS stimulation-online design applied in our study) (Figure 1).

**Figure 1.**
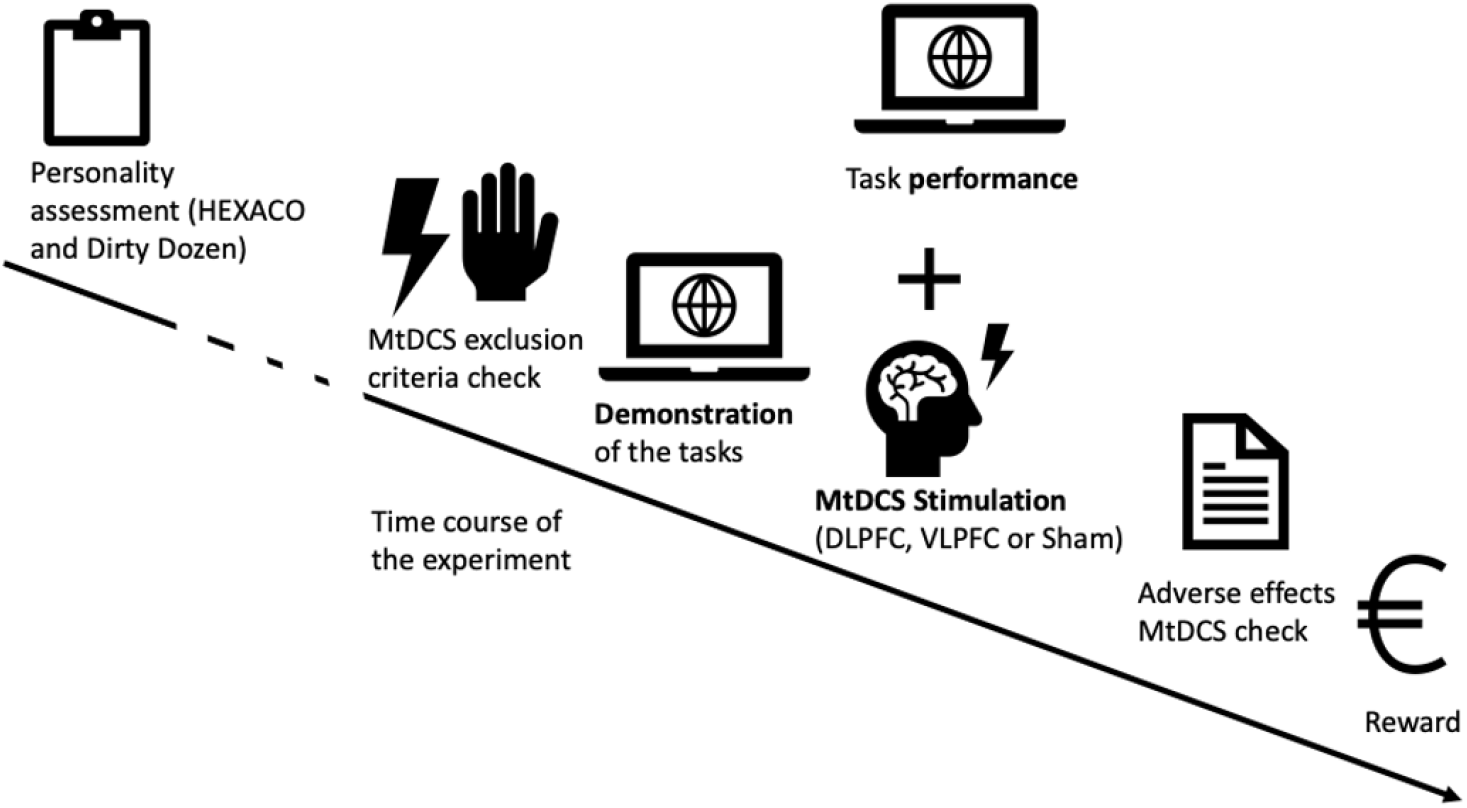
Representation of the experimental procedure. Prior to the experimental sessions, participants underwent a personality assessment (HEXACO-60 and dark-triad/dirty-dozen instruments), were checked against the exclusion criteria, and had the tasks demonstrated to them. Participants followed the same procedure during the different stimulations (DLPFC, VLPFC, and sham), held one day apart. Risk decision-making task (autoBART) order was randomized for all sessions and participants. The average time for both tasks combined was 10 minutes. The conduction time of each experimental session was 50 minutes: first 10 minutes we informed participants of the procedure and rated their mood pre-experiment, 20 minutes of stimulation (inside those 20 minutes the last 10 minutes participants performed both tasks) and the last 10 minutes consisted on post-experiment mood and adverse HD-tDCS effects assessment and financial compensation (last session).

Before the first HD-tDCS session, participants completed 2 questionnaires (see below) to profile their personality post-hoc; we also informed participants that the more points they accrued during task performance, the better they would be compensated. At the end of each session, participants rated their mood and pain/discomfort on a 4-point visual analogue scale, with zero participants rating any discomfort or pain during all sessions.

### 2.3 Randomization and blinding

To ensure the random and counterbalanced order of all stimulation sessions, a randomized list, created via an ad hoc web tool, determined the stimulation order of the three sessions to be followed by each participant. The study design and the specific stimulation condition delivered on each of the three sessions were blinded to participants, who were kept unaware of the type of stimulation they received (sham or active HD-tDCS).

### 2.4. Personality questionnaires

Personality questionnaires administered via the Qualtrics platform to all participants prior to their participation in the study were used to re-analyse the main dataset of our study taking into account personality profiles.

Psychologists distinguish between broad personality dimensions and narrow personality dimensions. The former summarize a large amount of information and are predictive of an important number of outcomes (Soto and John, 2017; Soto, 2019), although to a limited extent, a phenomenon known as ‘broad bandwidth’. In contrast, narrow personality dimensions have more precise definitions of behaviour, and they are likely to predict less behaviours, although with greater accuracy (Paunonen and Ashton, 2001). All together this phenomenon is dubbed ‘fidelity’. As in the physics domain, broad bandwidth is usually associated with low ‘fidelity’ and the other way around (Ones and Viswesvaran, 1996)

In order to achieve a trade-off between ‘bandwidth’ and ‘fidelity’, the two following personality questionnaires, the HEXACO-60 and the *Dark triad-dirty dozen*, were selected:

*The HEXACO-60* assesses broader personality dimensions. It consists of 60 questions assessing 6 domains, each composed of 10 items (Ashton and Lee, 2009), as follows: honesty-humility, emotionality, extraversion, agreeableness, consciousness, and openness to experience. Psychometric properties, including levels of internal consistency, inter-item correlations and test-retest reliability have been all properly validated (Ashton and Lee, 2009; Roncero *et al*., 2014). With the HEXACO, a greater perception of risk associates with emotionality, while greater conscientiousness links to fewer perceived benefits of risk (Weller and Thulin, 2012). Honesty-humility is associated positively with non-gambling propensities (McGrath *et al*., 2018), while conscientiousness reported to be the strongest positive predictor of decision-making performance (Weller *et al*., 2018).

*The dark triad-dirty dozen* is a 12-item personality inventory that simultaneously assesses the 3 dark-triad traits associated with personality: Machiavellianism (e.g., “I have used deception or lying to get what I want”), psychopathy (e.g., “I tend to have no remorse”), and narcissism (e.g., “I tend to want others to admire me”). This inventory has demonstrated, despite its brevity, excellent psychometric properties, more than adequate temporal stability and internal consistency, and excellent validity (Jonason and Webster, 2010). Interestingly, research linked a high degree of self-reported psychopathy to more risk-taking in the BART (Hunt *et al*., 2005).

### 2.5. High-Definition transcranial direct current stimulation

A STARSTIM 8 5G wireless hybrid EEG/tCS 8-channel system (NE, Neuroelectrics, Barcelona, Spain), with a constant current DC neurostimulator and 6 12-mm Ag/AgCl sintered electrodes (NG Pistim) were used to deliver tDCS stimulation. The contact area for the electrodes was π cm^2^.

To modulate the left VLPFC excitability, a cathode was placed on scalp site F7 according to the international 10-20 electroencephalogram (EEG) system (Jurcak *et al*., 2007), while 5 (return) electrodes were positioned in FP1, F3, FC5, FT7, and F9 locations around the former. To target the left DLPFC, a cathode was placed on F3, while 5 other (return) electrodes were located on AF3, FC1, FC3, FC5, and F5 (Nikolin *et al*., 2015; Guo *et al*., 2018) (Figure 2). We selected cathodal stimulation due to its well-established modulatory effects on causal studies and unilateral left prefrontal to avoid the difficulty to assign effects to either cathodal or anodal stimulation.

**Figure 2.**
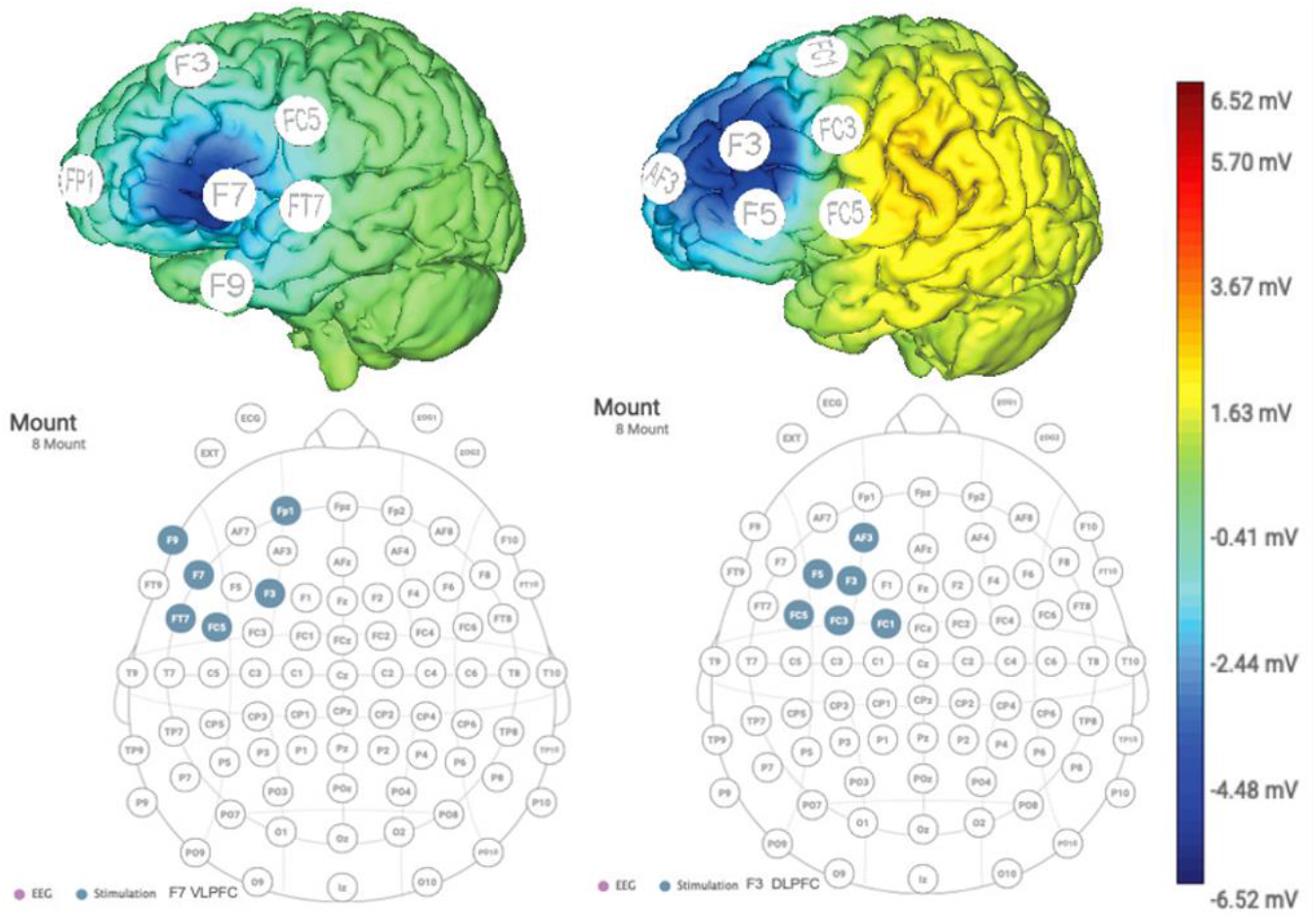
HD-tDCS montage. Six NG Pistim electrodes were placed as follows: left DLPFC (cathodal F3, return AF3, FC1, FC3, FC5, and F5), and left VLPFC (cathodal F7, return FP1, F3, FC5, FT7 and F9). The upper part of the figure shows computational models of the MtDCS montages used. The lowest voltage magnitude is shown at the approximate location of the cathodes (F3 and F7). The scale bar on the right shows the color codes for current intensity values (mV). The voltage distribution is shown on the realistic head model included in NIC 2 software (Neuroelectrics, Barcelona, Spain), which is based on the Colin27 dataset. Methods for generation of this head model and for electric field calculation can be found in (Miranda *et al*., 2013).

Once the neoprene cap was in place, we separated the hair, and cleaned the scalp with alcohol underneath each electrode location. The fastener component was then filled with conductive gel (Sigma Gel, Parker Laboratories, New Jersey, USA) using a curved syringe. HD-tDCS current was applied for a total duration of 20 minutes including a current ramp up and ramp down of 30 seconds at the beginning and at the end of the sham stimulation blocks. Subjects started performing the risk-taking task after 10 minutes of HD-tDCS. Since no available research analysed potential parametric effects of current intensities on a risk decision-making task before, two potentially effective HD-tDCS intensities 1.5 mA (current destiny 0,47 mA/cm^2^) and 2 mA (current destiny 0,63 mA/cm^2^) were tested in separate groups and treated as an independent variable.

Sham stimulation, was identical to active cathodal HD-tDCS to the left DLPFC but with the tDCS field off during the stimulation sessions. Specifically with 30 seconds of ramp up at the beginning and 30 seconds of ramp down at the end. None of the participants reported any of the classically known side-effects (e.g., itching, pain, headache, etc.) during or following stimulation.

### 2.6. Risk decision-making task

Risk decision-making was measured with a well-established computer-based paradigm, the Balloon Analogue Risk Task (BART) which is a behavioural measure of risk-taking. Performance in this task correlates with scores on measures of sensation seeking, impulsivity, and deficiencies in behavioural constraint (Lejuez *et al*., 2002). The BART is a widely used tool which has been proven a useful in the assessment of risk-taking previously (Sela *et al*., 2012; Seaman *et al*., 2015; Petrova and Garcia-Retamero, 2016; Gilmore *et al*., 2017; Russo *et al*., 2017; Gilmore *et al*., 2018; Nejati *et al*., 2018). It has also shown convergent validity with real-world risk-related situations (Fecteau, Pascual-Leone, *et al*., 2007).

We used a modified version of the BART (Lejuez *et al*., 2002), known as the autoBART (Figure 3) which maintains BART’s validity as well as unbiased statistics (Pleskac *et al*., 2008), in which participants are presented with a computer game in where they are requested to inflate a series of 30 ‘virtual’ balloons (see Figure 3). Participants were informed beforehand that they would be able to pump 127 times each ‘virtual’ balloon and that each one had a probability of bursting, set to 1/128 for the first pump, 1/127 for the second pump and so on, until the balloon exploded. Participants indicated in a text box how many times they chose to inflate the balloon to a maximum of 127 pumps. For each pump, participants earned a point and if the balloon exploded, the score was set to zero for that trial. Previous studies have shown that the adjusted average number of pumps of unexploded balloons (mean number of pumps on trials that the balloon did not explode) (AVP) indicates greater risk-taking propensity (Lejuez *et al*., 2002; Aklin *et al*., 2005; Cheng *et al*., 2012; Guo *et al*., 2018). Hence, we calculated the AVP, the total earnings (TE, i.e., the total accumulation of points for non-exploded balloons across all trials), and wanted pumps (WP, i.e., the total number of times the participant wanted to pump the balloon across all trials) as dependent variables.

**Figure 3.**
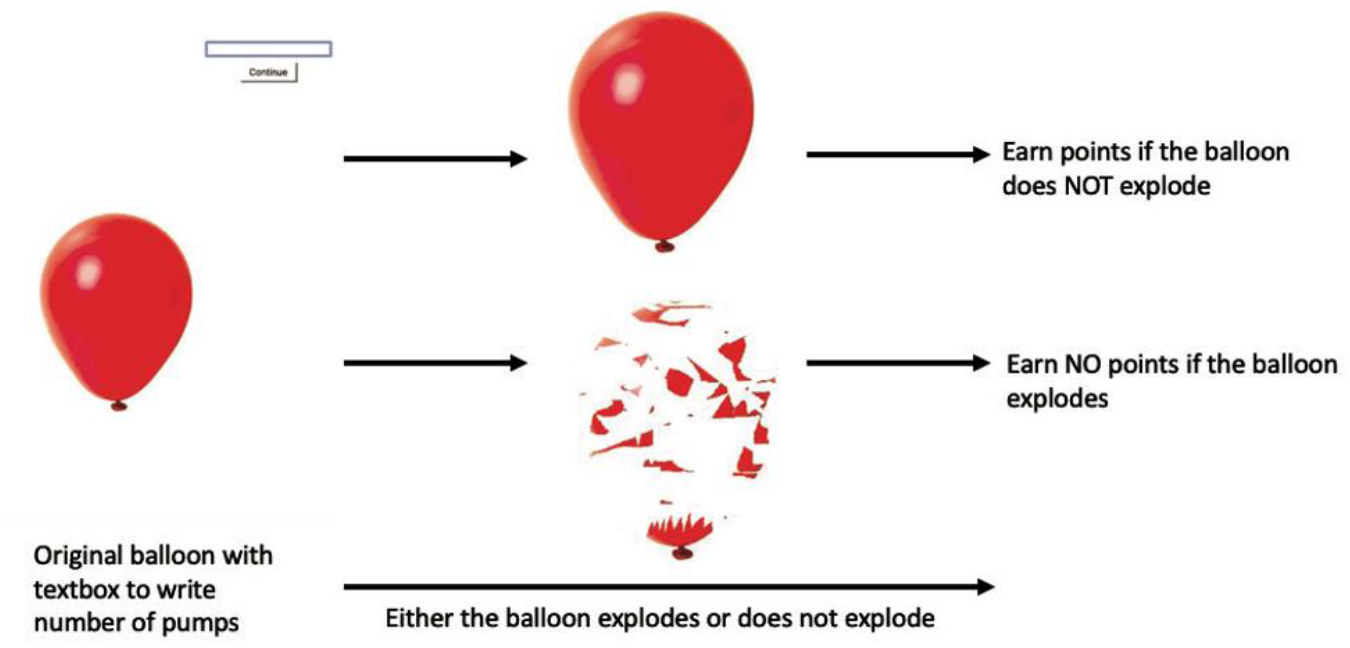
BART. Participants are instructed to achieve as many points as possible by inflating a series of 30 balloons but are warned of the probability of the balloon bursting. Shown a balloon and a text box, they are instructed to indicate how many times to inflate the balloon (maximum 127 times).

A point-to-euro conversion for the task was computed to motivate participants to accumulate as many points as possible and ultimately reach a higher reward. Furthermore, we programmed the task so the maximum point-to-euro conversion would be €30. The average financial compensation was €22. However, upon the request of the university’s ethics committee, at the end of the experiment, all participants had to be rewarded with the same financial compensation of €30.

### 2.7. Main analysis addressing the impact of left prefrontal HD-tDCS on risk taking

The analytical strategy of the study was designed pre hoc, as a single-blinded mixed factorial 3×2 design. Stimulation condition (DLPFC, VLPFC, sham), as within-subject factor, and stimulation intensity (1.5 mA or 2 mA) as between-subject factor. Post hoc, an additional between subject factor, personality (3×2×*n*), determined by a Latent Profile Analysis (LPA) was added in a subsequent re-analysis of the same dataset considering the influence of personality traits.

Main analyses were carried out considering independent variables as categorical, and dependent variables as continuous. For the first analysis, categorical variables were cathodal HD-tDCS target condition (3 levels: DLPFC, VLPFC, sham) and HD-tDCS stimulation intensity: (2 levels: 1.5 mA, 2 mA). The continuous variables were autoBART’s AVP, TE, and WP outcome measures. We subjected participants to repeated measures of cathodal stimulation (HD-tDCS) effects on risk taking. Mixed analyses of variance with the corresponding contrast analyses (simple for the between-group effect, and polynomial for the within-group effect) were performed. Likewise, the Shapiro-Wilk and Levene tests were used to verify we were analysing random samples from normal populations with the same variance, while the Mauchly test was used to test the sphericity assumption.

We calculated effect-size measurements (Cohen’s d and eta-squared-η^2^-) and its corresponding confidence intervals (CI) for a better understanding of the relative magnitude of the experimental changes. Following previous literature, to obtain a confidence interval that was equivalent to the ANOVA F test of the effect (which employs a one-tailed, upper tailed, probability) we applied a CI of 90% (Steiger, 2004; Clay, 2014; Lakens, 2014).

Additionally, given that the limited sample size of our study might potentially limit the statistical power, Bayesian analyses were performed to determine whether a non-significant effect truly signalled the lack of an intervention effect (Biel and Friedrich, 2018). We tested the relative plausibility of the alternative hypothesis (H_1_) over the null hypothesis (H_0_), i.e., the presence and absence, respectively of the effects of HD-tDCS target/condition and stimulation intensity on autoBART performance. We calculated the Bayes factor expressed as BF10, using the counterpart Bayesian tests of the analyses described above, for a credible interval of 95%. Since we had no previous data with which to establish an informed prior, the default Cauchy prior width of 0.707 provided by JASP 0.12.2 was used (JASP-Team, 2020). We compared the models used for analyses to the model containing the grand mean and the random factors, called the null model. Frequentist analyses were performed using SPSS version 23 (IBM Software Group, IL, USA) and STATA version 16 (StataCorp LLC, USA), while JASP computer software, version 0.12.2 (JASP Team, Amsterdam, The Netherlands) was used for Bayesian analyses.

### 2.8 Post hoc analyses to assess the influence of personality traits

We reanalysed the same dataset to assess a potential impact of personality profile on HD-tDCS stimulation effects delivered actively to the VLPFC or the DLPFC vs sham tDCS stimulation. To classify participants by personality profile, we conducted a latent profile analysis (LPA) of the personality data previously collected.

LPA recovers latent groups from observed data and clusters individuals into groups with similar characteristics in relation to a set of measured variables (Steinley and Brusco, 2011; Flaherty and Kiff, 2012; Oberski, 2016). It shows additional patterns of relationships above and beyond regression analyses (Stanley *et al*., 2017). We determined the number of participants in each class empirically, as there were no a priori assumptions regarding the number of individuals on each class. However, we computed some simulation experiments in order to estimate the minimum sample size required to achieve a power of 0.80 for LPA classification using different indicators.

Simulations showed that a sample of 30 participants was not enough to retrieve the correct number of profiles using either the Akaike Information Criterion or the Bayesian Information Criterion. These were outperformed, though, by the Sample-Adjusted Bayesian Information Criterion (SABIC), which was able to recover the correct number of classes even when the average effect size between classes was ∂ (Cohen’s d) = 0.2. Accordingly, we used the SABIC for the determination of the number of latent profiles of this study. All code, simulations, and results are available at: https://osf.io/m79pg/?view_only=887f5b53d3b14319898afac7b7391885. Finally, R Studio 1.1.463 and tidyLPA package (M. Rosenberg *et al*., 2018) was employed to conduct the LPA of the personality data.

## 3. RESULTS

### 3.1 Impact of prefrontal stimulation on risk taking behaviour

The Shapiro-Wilk test indicated that the dependent variables (autoBART: AVP *W*=0.310, *p*=0.735), TE *W=2.324, p*=0.115), and WP W=0.101, *p*=0.904)) were normally distributed in the 3 HD-tDCS conditions (DLPFC, VLPFC, and sham). The Levene’s test showed equality of variances for the same 3 conditions (AVP: DLPFC *p*=0.297, VLPFC *p*=0.529, Sham *p*=0.099; TE: DLPFC *p*=0.215, VLPFC *p*=0.060, Sham *p*=0.696; WP: DLPFC *p*=0.173, VLPFC *p*=0.242, Sham *p*=0.223). For the repeated measures’ analysis, the Mauchly test indicated that the variances of the differences between all possible pairs of within-subject conditions (HD-tDCS) were equal (sphericity was assumed) (TE *X^2^(2) = 4.419, p =0.110*, WP *X^2^(2) = 3.362, p = 0.186*, and AVP *X^2^(2) = 1.292, p = 0.524*).

No significant main effect of stimulation intensity on the dependent variables (auto BART: AVP *F*_(2,31)_=0.20, *p*=0.735, TE *F*_(2,31)_=2.324, *p*=0.115, and WP *F*_(2,31)_=0.101, *p*=0.904) was observed, for any HD-tDCS condition, suggesting that differences in this parameter did not have any bearing on the magnitude of the modulatory effects generated by HD-tDCS on risk-taking during the autoBART trials (See table 1 for the detailed descriptive analysis of the dependent variables).

**Table 1.**
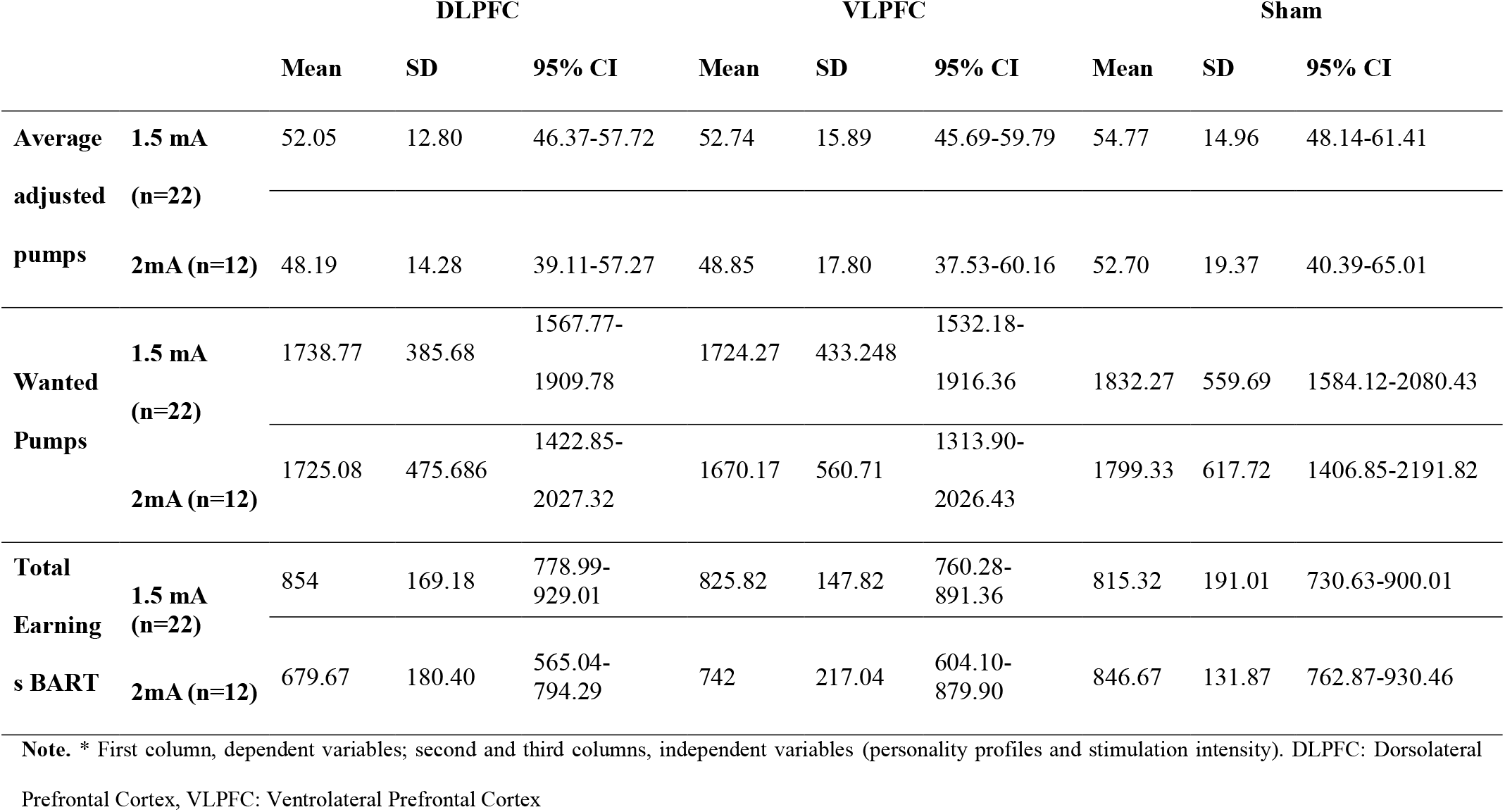
Descriptive analysis of dependent variables.

In relation to the AVP dependent variable, which was the most directly related with risk-taking, a mixed-design analysis of variance showed a significant main effect (within-group effect) of the stimulation condition (MtDCS: *F*_(2,64)_=3.612, *p*=0.033; observed power =0.648; η^2^=0.101, 90% CI [0.000-0.2160]; polynomial contrast *F*_(1,32)_=7.770, *p*=0.009; observed power =0.771; η^2^=0.195, 90% CI [0.0354-0.4101]). Specifically, the planned post-hoc paired *t*-test (Bonferroni adjustment for multiple comparisons) revealed a lower AVP (hence lower risk-taking) under DLPFC stimulation vs sham (*t* _(31)_=2.32, p=0.027; mean difference (Mdiff) =3.619; Cohen’s d= 0.57, 95% CI [0.71-1.07]) (See Figure 4).

**Figure 4.**
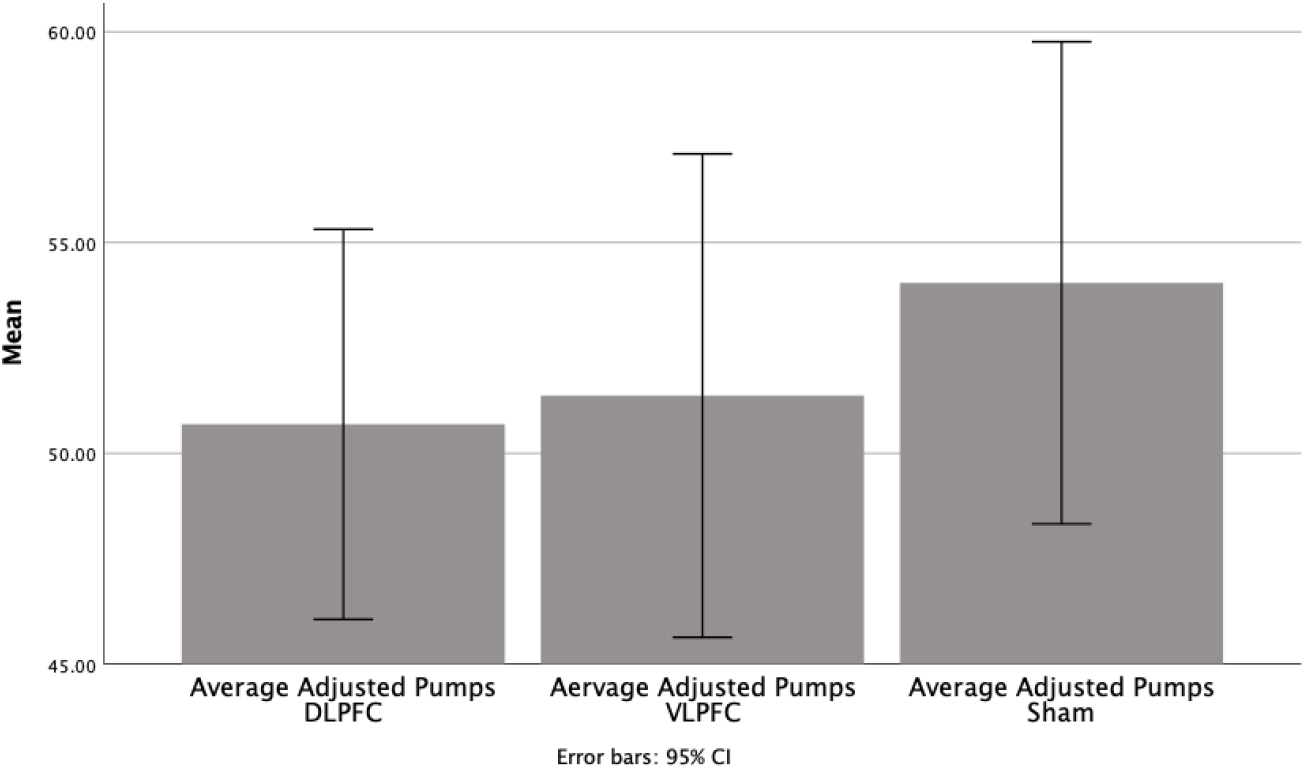
Adjusted average of pumps (AVP) in the autoBART task. Comparison of the means of the tDCS conditions (DLPFC, VLPFC and Sham). The figure represents the arithmetic means of each group plotting the 95% confidence interval (CI) of the mean. DLPFC: Dorsolateral Prefrontal Cortex, VLPFC: Ventrolateral Prefrontal Cortex.

A mixed-design analysis of variance showed a pattern toward significance regarding the differences in WP for the within-group effect (MtDCS: polynomial contrast *F* _(2,64)_=2.954, *p*=0.059; observed power =0.556; η^2^=0.085, 90% CI [0.000-0.309]). (See Figure 5 and 6).

**Figure 5.**
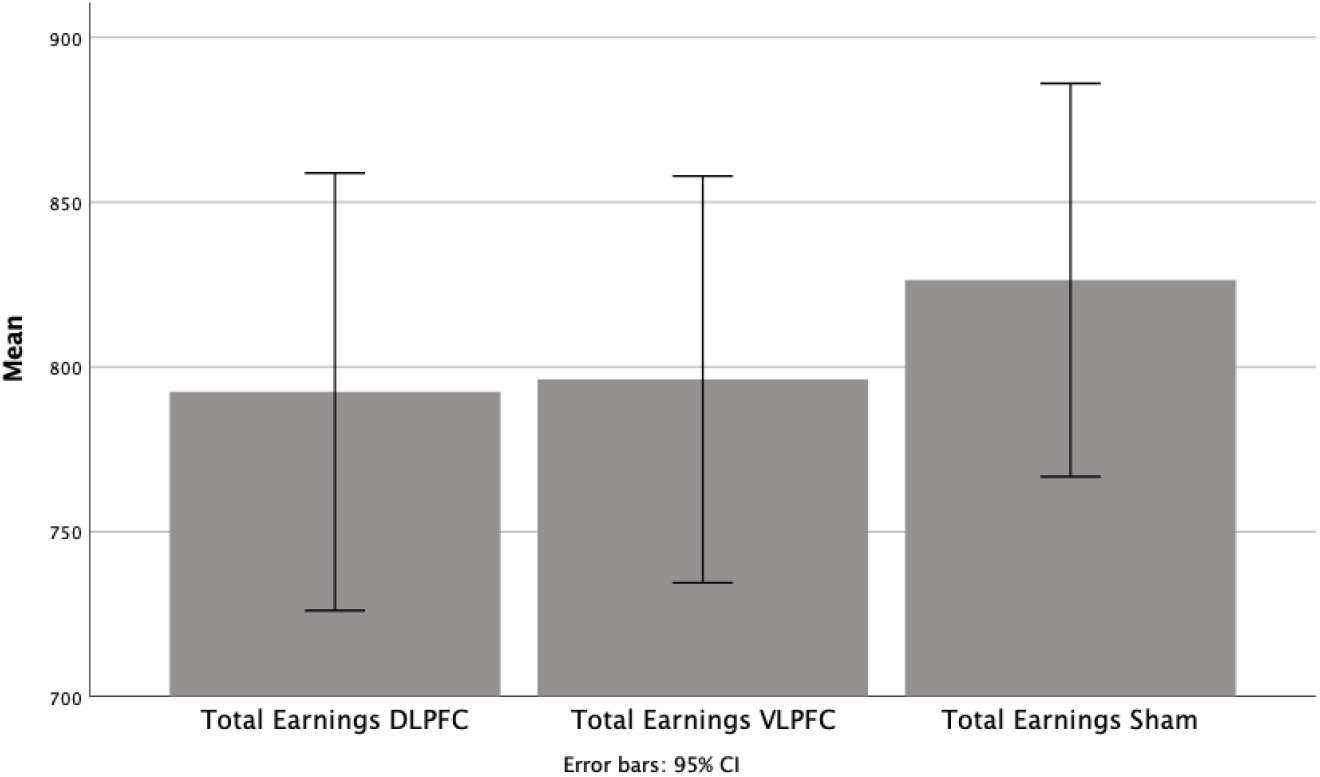
Total earnings (TE) in the autoBART task. Comparison of the means of the tDCS conditions (DLPFC, VLPFC, Sham). The figure represents the arithmetic means of each group plotting the 95% confidence interval (CI) of the mean. DLPFC: Dorsolateral Prefrontal Cortex, VLPFC: Ventrolateral Prefrontal Cortex.

**Figure 6.**
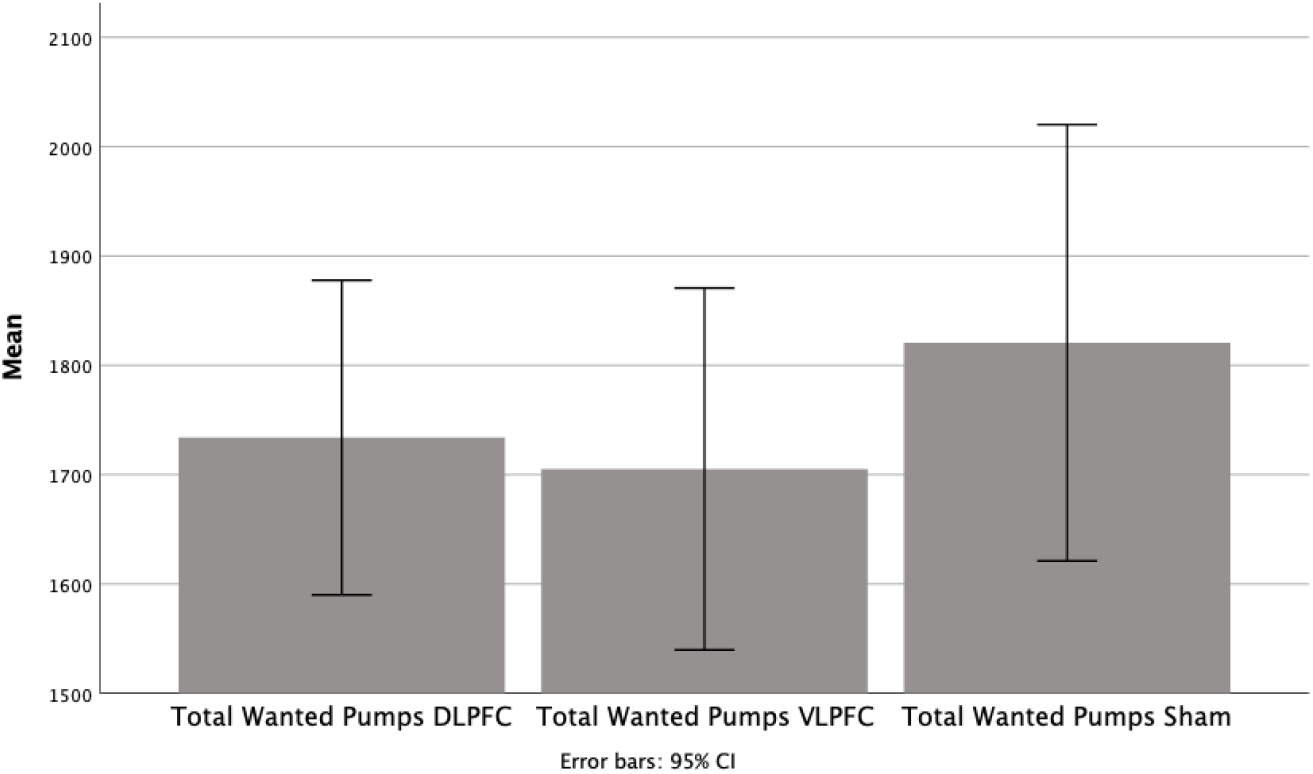
Wanted pumps (WP) in the autoBART task. Comparison of the means of the three tDCS conditions (DLPFC, VLPFC and Sham). The figure represents the arithmetic means of each group plotting the 95% confidence interval (CI) of the mean. DLPFC: Dorsolateral Prefrontal Cortex, VLPFC: Ventrolateral Prefrontal Cortex.

### 3.2 Influence of personality traits on the tDCS modulation of risk-taking behaviour

As indicated in the methods section, we performed a re-analysis of our HD-tDCS DLPFC and VLPFC and sham stimulation datasets reported in the prior section, taking into consideration personality profiles (See Supplementary Materials Table S1 for descriptive analysis of personality data).

To this end we extracted personality features, from that original sample of participants collected prior to participation in their 1st HD-tDCS sessions and assessed the best-fitted profile, by examining 3 models and selected a 3-class model comparing interpretability and statistical soundness (Sample Adjusted-Bayesian information criterion (SABIC) 1795.729). We compared the former profile with 2-profile and 4-profile models of higher BIC and lower Entropy (Table 2). The 3-profile latent model exhibited the best trade-off between SABIC, BIC, and Entropy^1^ (Vrieze, 2012; Criterion, 2015; Stanley *et al*., 2017; Araújo *et al*., 2019) and was retained for subsequent analyses.

**Table 2.**
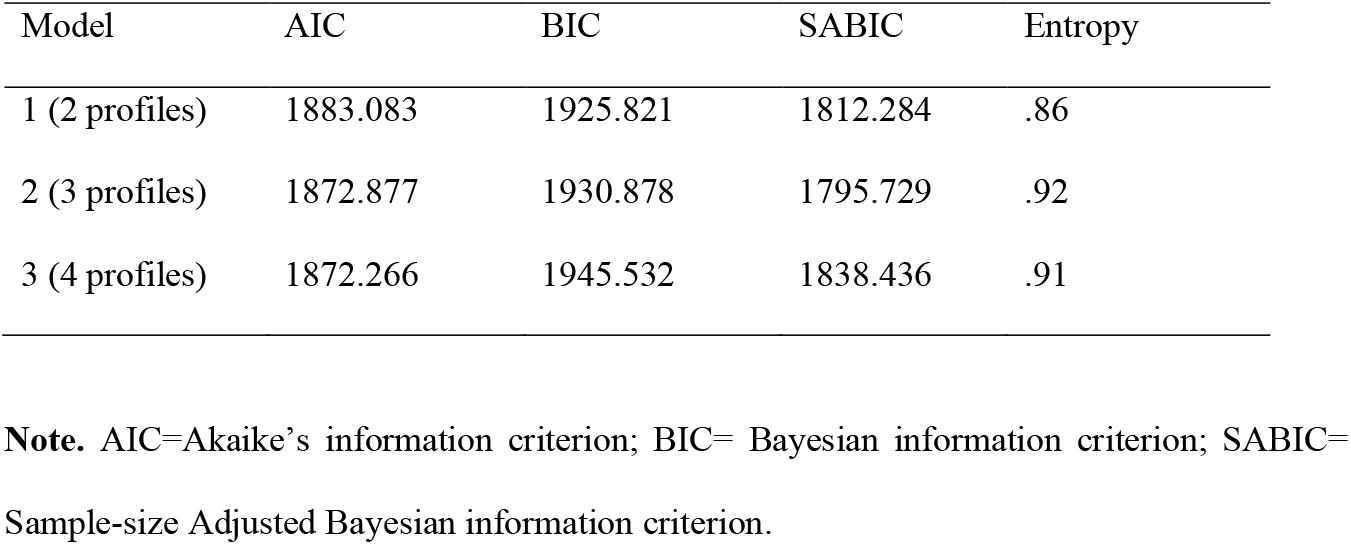
Results for competing latent profile analysis models of personality data.

Profile 1 (n=10) was consistent with dark-triad behaviours (Lee and Ashton, 2014), and thus, scored high on Machiavellianism, narcissism, psychopathy, and extraversion, and low on agreeableness, conscientiousness, emotionality, honesty-humility, and openness to experience. Therefore, was labelled as the *impulsive-disinhibited* group. Profile 2 (n=18) reflected a pattern of centred, homogenous means, hence was labelled the *normative* group. Finally, profile 3 (n=6), which scored around the mean in dark-triad behaviours, low on extraversion and agreeableness, and high on emotionality, was labelled the *inhibited/emotional* group, and was to be expected as lower risk-taking (see Figure 7 for a graphic representation) than the other two. Table 3 shows average values means for the dimensions for each personality profile, identifying 3 profiles for interpretation and labelling. Correlations between dependent variables and personality dimensions are reported in Table S.2 in the Supplementary Materials.

**Figure 7.**
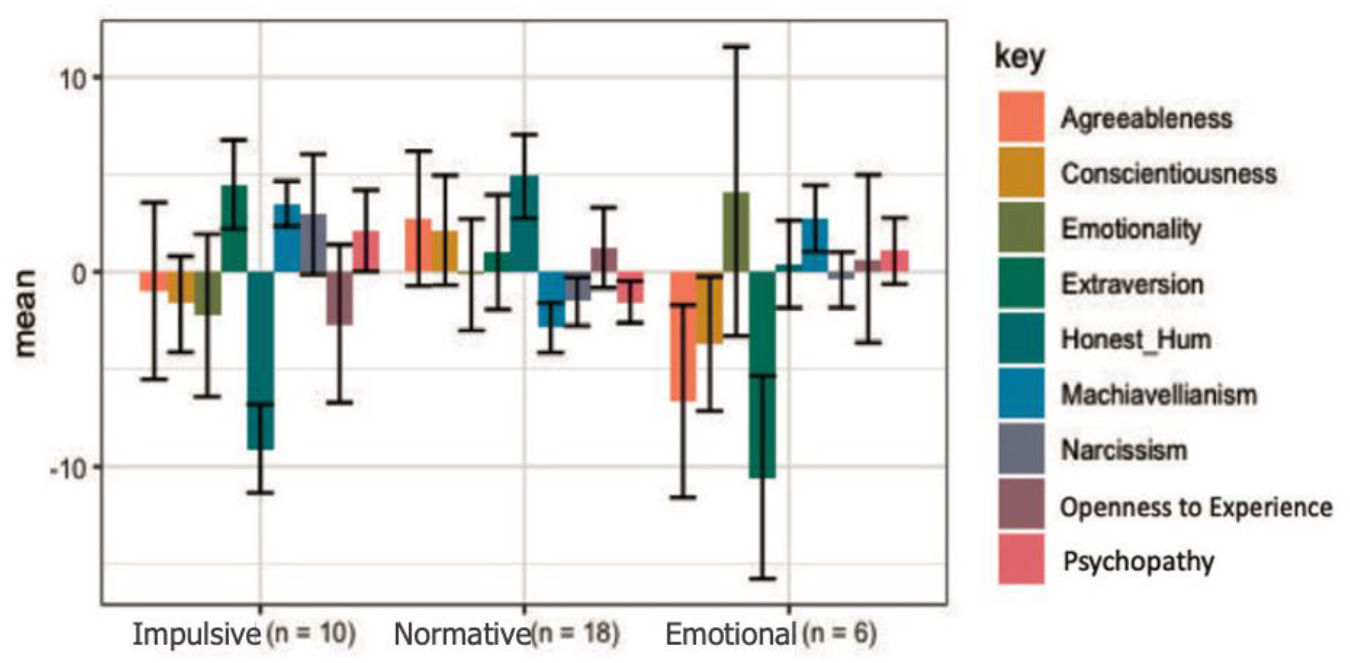
Personality profile estimation. Personality profile model specified by passing arguments to the *variance* and *covariance* arguments. In this model the equal variances and covariances are fixed to 0 by default.

**Table 3.**
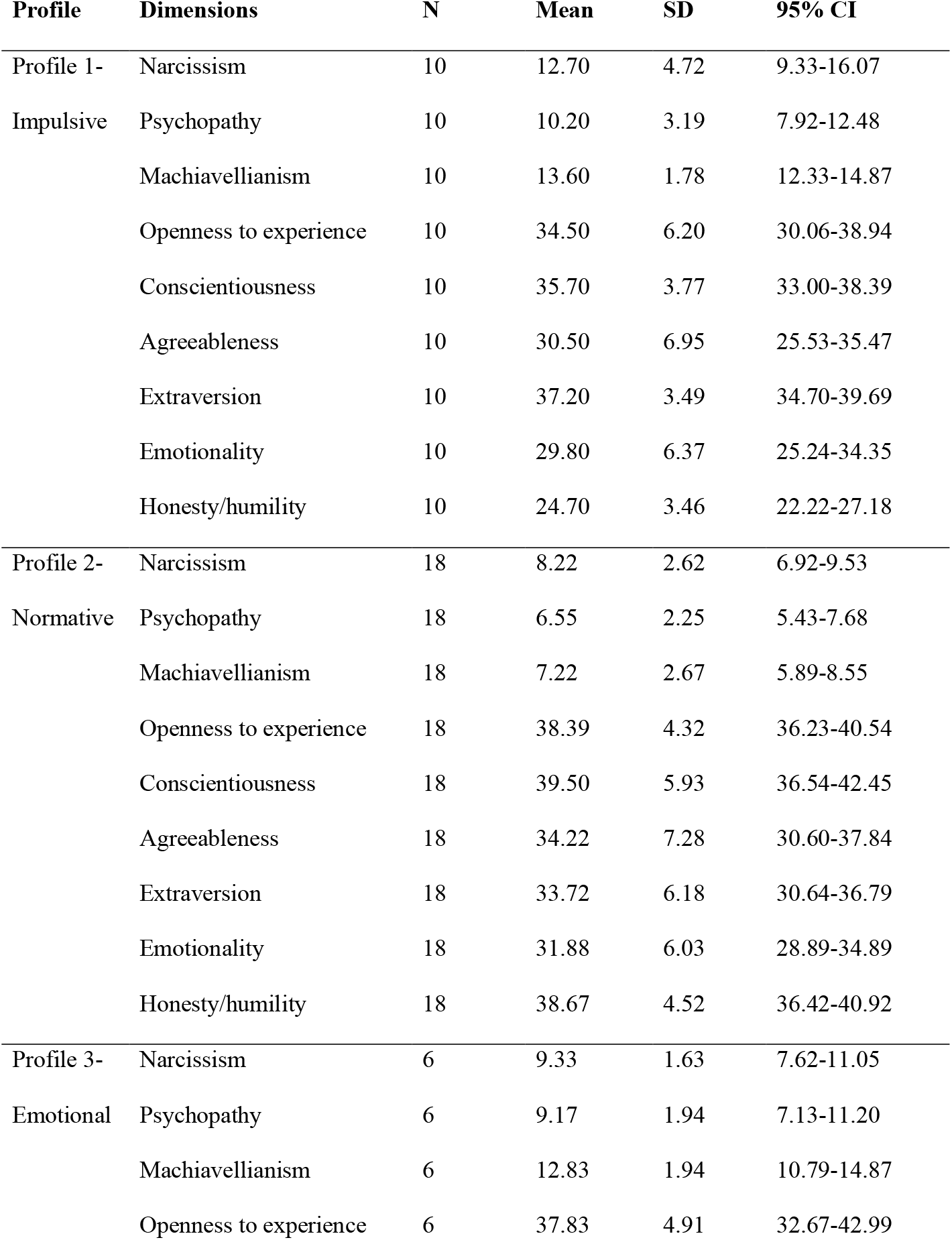
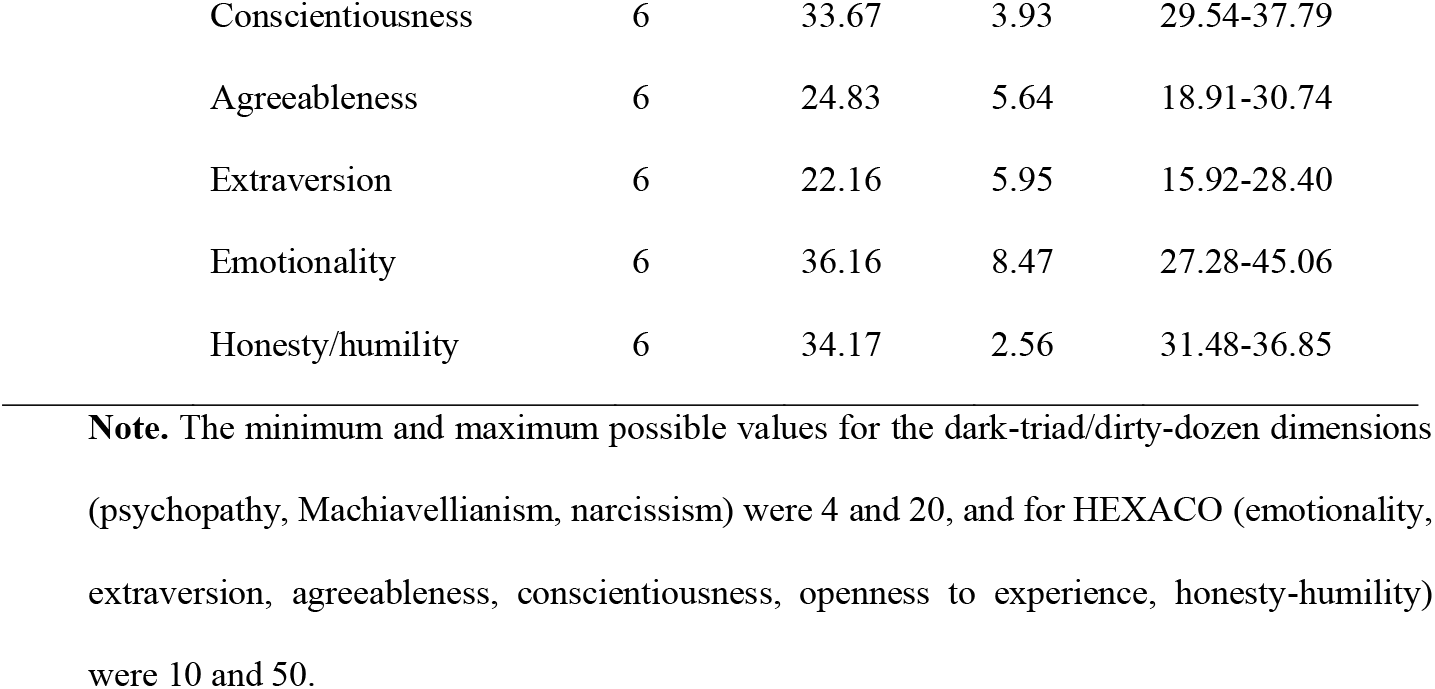
Descriptive analysis of personality dimensions by profile.

Although the interaction between personality and stimulation condition did not reach statistical significance for the autoBART variable AVP (Adjusted Average Pumps for non-exploded balloons) for personality profile 1, a pattern toward significance regarding differences between the three HD-tDCS conditions was identified (*F*_(2,27)_=3.142, *p*=0.059; observed power =0.554; η^2^=0.1888 90%, CI [0.000-0.3555]). Specifically, a post-hoc paired *t*-test (Bonferroni adjustment for multiple comparisons) revealed lower AVP for DLPFC stimulation vs sham (*t* _(27)_ =2.05, *p*=0.0502; Mdiff=9.305; Cohen’s d= 0.497 95%, CI[-0.00-0.99]), indicating that profile 1 under DLPFC cathodal stimulation showed decreased risk-taking vs sham. For personality profile 2, despite the fact that the analysis of variance showed significant differences between the 3 HD-tDCS conditions (*F_(2,27)_*=3.754, *p*=0.036; observed power =0.636; η^2^=0.217, 90% CI [0.008-0.384]), post-hoc paired *t*-test (Bonferroni adjustment for multiple comparisons) did not point to significant differences for any of the comparisons (see figure 8).

**Figure 8.**
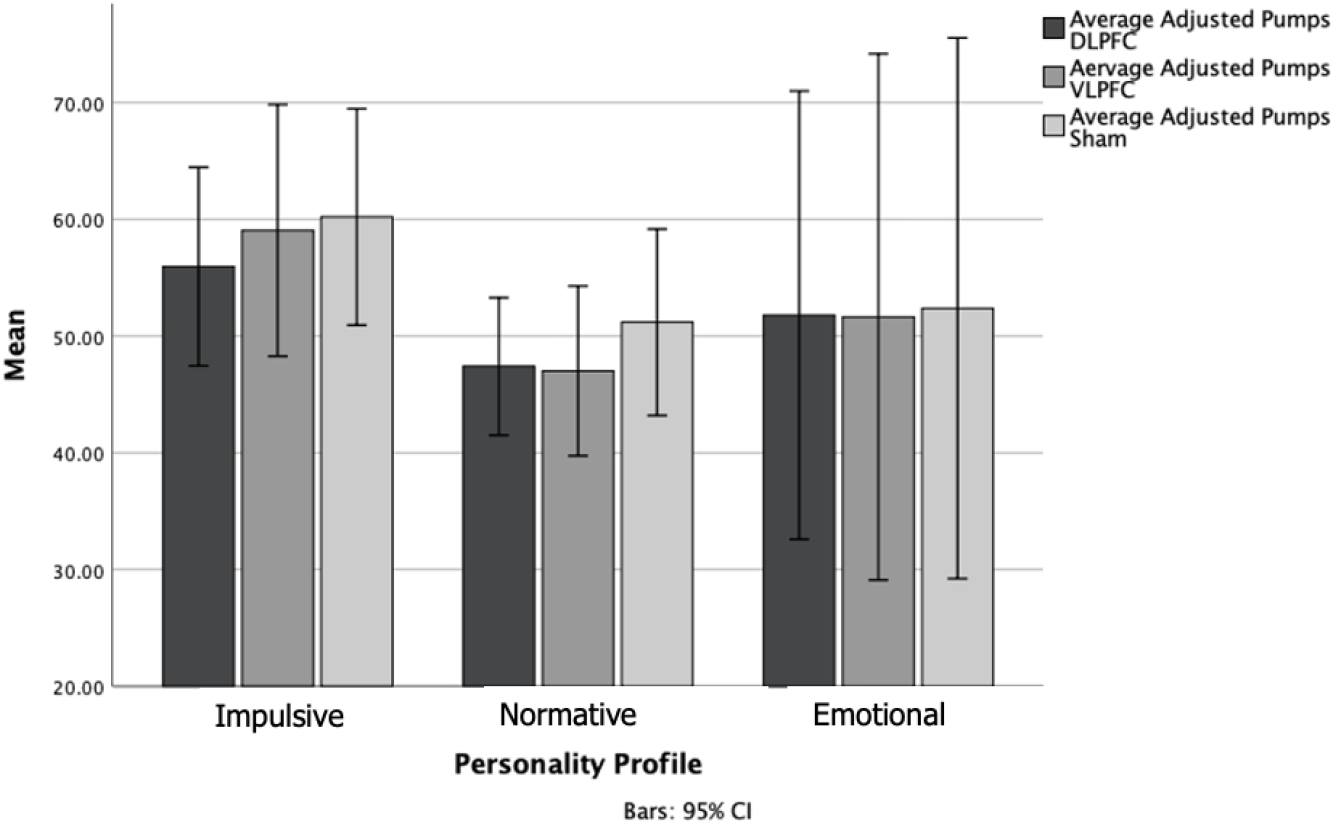
Graphic distribution of adjusted average of pumps (AVP) in the autoBART task. Comparison of the means of the three stimulation conditions (DLPFC, VLPFC and Sham) by personality profiles. The figure represents the arithmetic means of each group plotting the 95% confidence interval (CI) of the mean. DLPFC: Dorsolateral Prefrontal Cortex, VLPFC: Ventrolateral Prefrontal Cortex.

Regarding autoBART TE (Total Earnings), although a mixed-design analysis of variance showed no significant main effect for HD-tDCS (within-group effect), it did show a significant main effect for personality (between-group effect) (*F* _(2,28)_=6.155, *p*=0.0061; observed power =0.854; η^2^=0.3054, 90% CI [0.0615-0.4642]). The Bonferroni corrected post-hoc test showed higher TE values for personality profile 1 than for personality profile 2 (*t*_(28)_=2.59, p=0.0151; Mdiff=168.211; Cohen’s d= 1.02, 95% CI[0.19-1.83]), suggesting that profile 1 was more risk-taking.

A significant Interaction between the stimulation condition and personality, suggested that the effect of the tDCS on TE depended on the personality profile (HD-tDCS x personality: *F* _(2,28)_=3.524, *p*=0.043; observed power=0.608; η^2^=0.2011, 90% CI [0.0036-0.3660]). Specifically, for VLPFC stimulation, we found significant differences between the three personality profiles (*F*_(2,28)_=7.708, *p*=0.0022; observed power=0.925; η^2^=.3551, 90% CI [0.0985-0.5073]). Indeed, the Bonferroni corrected post-hoc tests revealed that the modulation of VLPFC by HD-tDCS, produced lower TE values for personality profile 2 than for personality profiles 1 and 3 (*t*_(28)_=2.56, *p*=0.0162; Mdiff=261.567; Cohen’s d= 1.01, 95% CI[0.19-1.82] and *t*_(28)_=2.62, *p*=0.0139; Mdiff= 217.567; Cohen’s d= 1.24, 95% CI[0.25-2.20] respectively). This result suggests lower risk-taking for profile 2 under VLPC stimulation (See figure 9)

**Figure 9.**
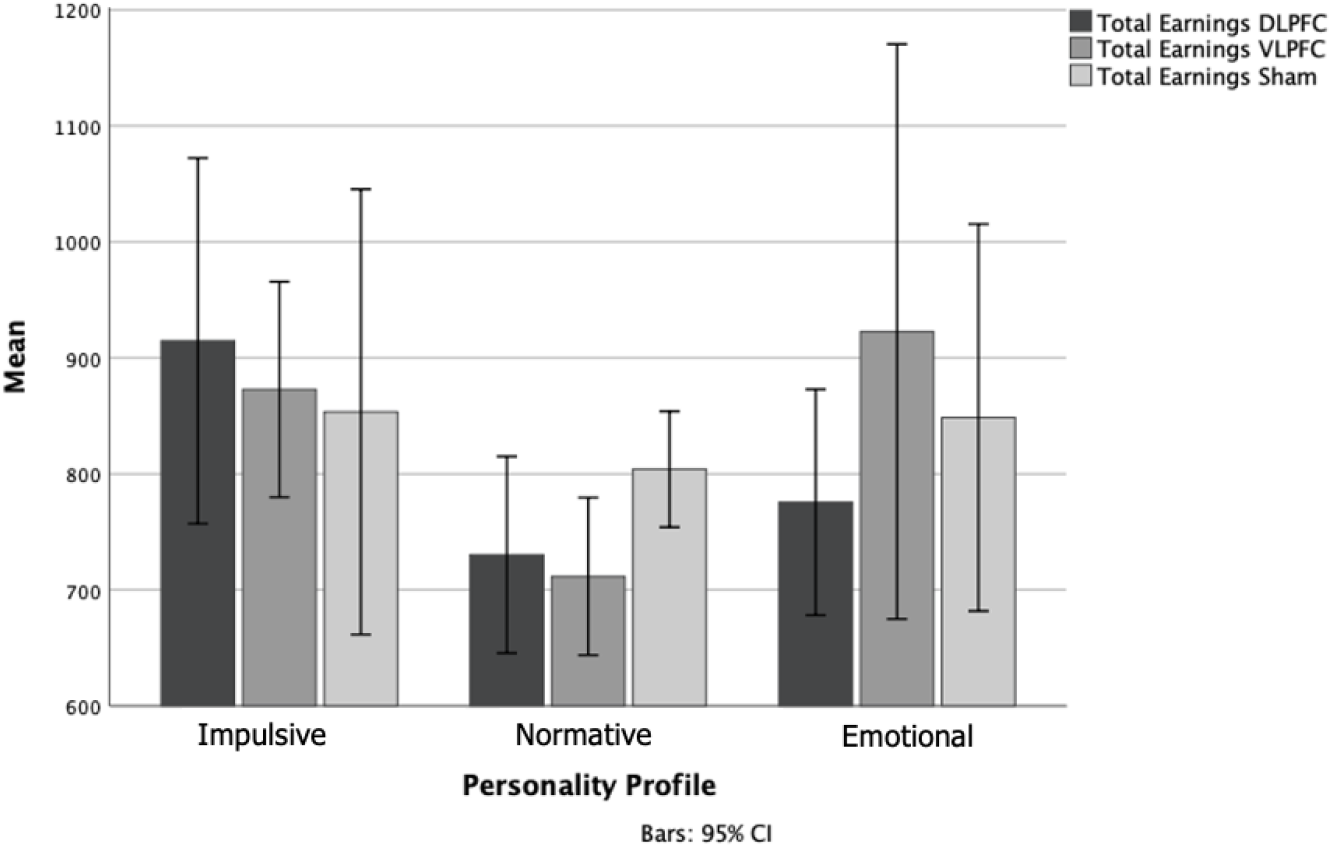
Graphic distribution of total earnings (TE) in the autoBART task. Comparison of the means of the three tDCS conditions (DLPFC, VLPFC and Sham) by personality profiles. The figure represents the arithmetic means of each group plotting the 95% confidence interval (CI) of the mean. DLPFC: Dorsolateral Prefrontal Cortex, VLPFC: Ventrolateral Prefrontal Cortex.

Although the analysis of variance yielded no significant main effect for personality (between-group effect) for the autoBART variable WP (Wanted Pumps), significant differences between the three HD-tDCS conditions were found for personality profile 1 (*F*_(2,27)_=3.598, *p*=0.0412; observed power =0.616; η^2^=0.2104, 90% CI [0.0049-0.3774]). Specifically, the Bonferroni corrected post-hoc test showed lower WP values for DLPFC stimulation vs sham (*t* _(27)_=1.99, *p*=0.0568; Mdiff=322.278; Cohen’s d= 0.48, 95% CI[-0.013-0.97]). Meaning that under DLPFC stimulation, this personality profile was less risk-taking (see Figure 10). For a pairwise comparison of cathodal DLPFC vs sham, cathodal VLPFC vs sham and cathodal VLPFC vs cathodal DLPFC under different personality groups see Supplementary Materials Table S3.

**Figure 10.**
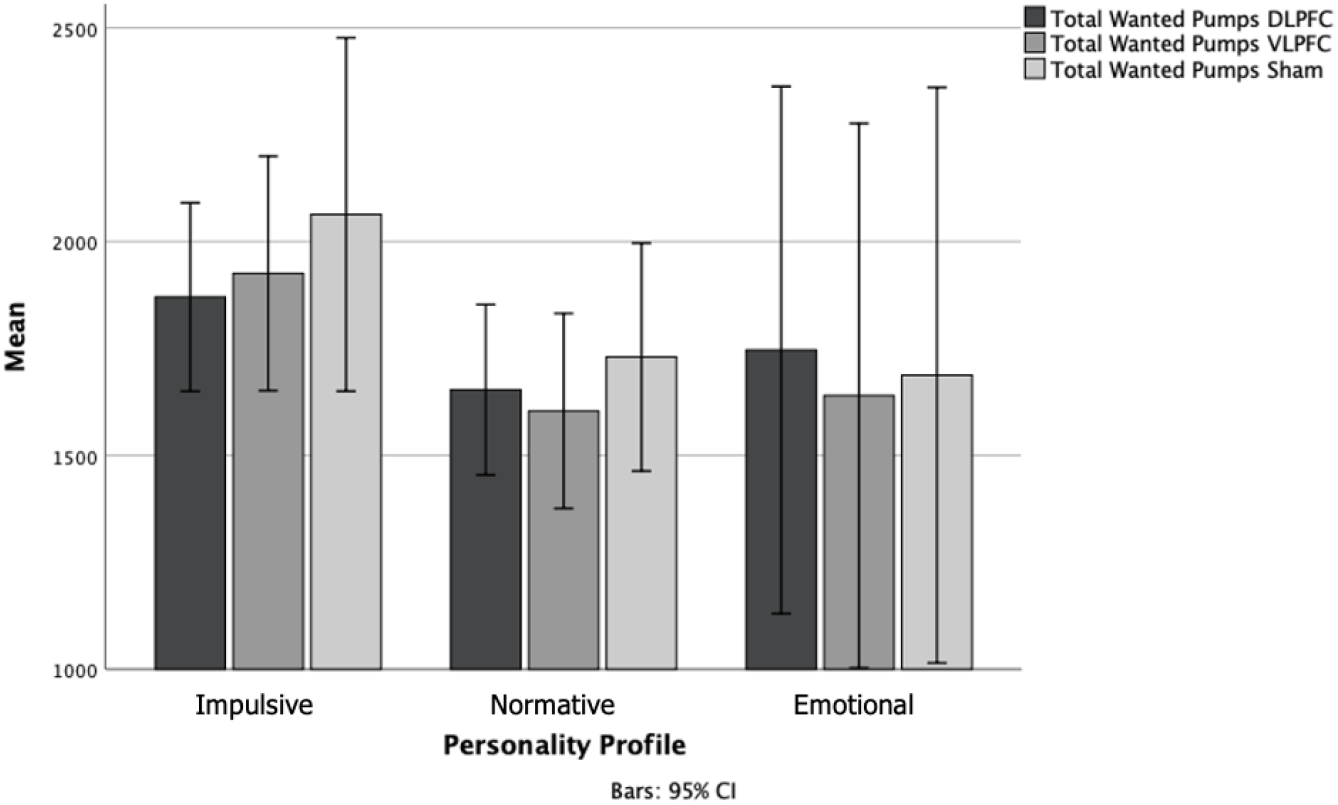
Graphic distribution of wanted pumps (WP) in the autoBART task. Comparison of the means of the three tDCS conditions (DLPFC, VLPFC and Sham) by personality profiles. The figure represents the arithmetic means of each group plotting the 95% confidence interval (CI) of the mean. DLPFC: Dorsolateral Prefrontal Cortex, VLPFC: Ventrolateral Prefrontal Cortex.

Comparisons using Bayesian methods between independent groups (personality and stimulation intensity) for the HD-tDCS condition (within-subject factors) and for each of the dependent variables yielded very similar results to those revealed by frequentist statistical analyses. Overall, the Bayesian analyses reinforced the finding of non-significant effects for the frequentist analyses (i.e., stimulation intensity) and support the null hypothesis.

Correlation analyses showed that psychopathy was positively and significantly correlated with AVP for all 3 MtDCS conditions (DLPFC: r=0.47, p=0.01; VLPFC: r=0.45, p=0.01; sham: r=0.44, p=0.01). Emotionality negatively correlated with all three HD-tDCS conditions (DLPFC: r=-0.41, p= 0.05; VLPFC: r=-0.44, p=0.01; sham: r=-041, p=0.05).

## 4. DISCUSSION

We here aimed at investigating in humans the causal contributions of the left DLPFC and the left VLPFC regions to risk decision-making operations. To this end, we carried out a neurostimulation study based on a 3-way comparison of two active cathodal stimulation conditions (DLPFC and VLPFC) and sham tDCS. We used HD-tDCS since the studied areas (VLPFC and DLPFC) were wide, hence very focal techniques such as TMS are less suited when the exact MNI (Montreal Neurological Institute) xyz coordinates of key cortical target are not necessarily clear. Secondarily, we took advantage of our study to reanalyse the same dataset and explored post-hoc, whether personality traits influenced the impact of prefrontal stimulation and the modulation of risk decision-making, initially described as main outcome of the study.

### 4.1 Dorsal and ventral dorsolateral prefrontal HD-tDCS on risk-taking behaviour

The autoBART used in our study signals lower less risk-taking behaviour in case of fewer wanted pumps (WP) and average adjusted pumps (AVP) for non-exploded balloons. In such context, our study showed that cathodal left DLPFC decreased both WP and AVP compared to sham and left VLPFC stimulation at both of the HD-tDCS intensities (1.5 and 2 mA) employed. Our study also revealed a marginal effect for the VLPFC on autoBART measures which did not ultimately reach significance. Hence, we conclude that cathodal stimulation of the left DLPFC led to a decrease of risk-taking behaviour. Additionally, our study, which is the first exploring the causal role of the left VLPFC in a risk decision-making via the autoBART, failed to find effects for the modulation of this ventral prefrontal area. Also importantly, the HD-tDCS-induced effects on risk decision-making where invariant to stimulation intensity -at least for the two different levels tested in our study-, an outcome in agreement with prior evidence from reporting no effect differences between these same intensities (Ho *et al*., 2016; Jamil *et al*., 2017).

Overall, the findings of our main analysis converge with prior reports in suggesting that either bilateral DLPFC cathodal tDCS stimulation (Fecteau, Pascual-Leone, *et al*., 2007; Fecteau, Knoch, *et al*., 2007; Paulo S Boggio *et al*., 2010; Minati *et al*., 2012; Cheng and Lee, 2016; Ye *et al*., 2016; Huang *et al*., 2017; Ota *et al*., 2019) or also unilateral left cathodal DLPFC stimulation (Sela *et al*., 2012; Guo *et al*., 2018) have the ability to foster lower levels of risk propensity and lower risk-taking.

In absence of complementary neuroimaging or neurophysiological evidence, both beyond the scope of our pilot study, it is risky to speculate on the potential networks and coding mechanisms involved in the above-reported outcomes. Previous studies have found that the number of balloon pumps in the BART task correlates negatively with whole-brain connectivity estimates for the right DLPFC, an outcome suggesting that a tDCS regime applied to this region may indirectly affect networks implicated in valuation and choice (Weber *et al*., 2014). Additionally, the neural bases, underlying risk decision-making, include different portions of the prefrontal cortex, encompassing lateral and medial areas. However, the lack of effects observed during the modulation of the left VLPFC targeted in our study (Fellows and Farah, 2007; Camus *et al*., 2009; Peter N C Mohr *et al*., 2010) suggests that this other prefrontal region might play a differential role in risk decision-making and supports the idea of two dissociable anatomical systems in the prefrontal cortex (see below for further discussion).

### 4.2 Personality and prefrontal tDCS modulatory effects on risk-taking behaviours

Prior studies have addressed the role of personality, especially extraversion and neuroticism in decision making, and assessed their impact on tDCS stimulation effects. For example, when modulated by tDCS, individuals with higher scores of personality dimensions related to introversion had a more evident effect compared to individuals with personality dimensions for extroverts (Peña-Gómez *et al*., 2011). On such basis, we re-examined our dataset and studied post-hoc if personality traits extracted from our population of participants could have affected the above reported impact of HD-tDCS stimulation or lack thereof in prefrontal regions.

Our analyses revealed that modulatory effects were more pronounced for *impulsive-disinhibited* participants and that *normative* participants tended to be less risk-taking in the autoBART. As a result, the latter earned less money (TE) than the former. Importantly, this effect interacted with the stimulated prefrontal region. This is why, participants in the *normative* group showed a significant pattern towards lower risk-taking behaviour under VLPFC stimulation compared to *impulsive-disinhibited* or *inhibited/emotional* participants. This outcome suggests that novel significant modulatory effects might emerge when behavioural datasets are specifically re-analysed taking into consideration the personality traits of participants.

The influence of personality profile on the ability of VLPFC to contribute to risk taking behaviour might also suggests the existence of two dissociable neural systems in the prefrontal cortex: a dorsal system encompassing the DLPFC specialized in cognitive regulation of risk decision-making, and a ventral system, including the VLPFC underpinning more focused on emotion on risk decision-making and the DLPFC (Yamasaki and Labar, 2002; Morawetz *et al*., 2020). To this regard, research has shown the involvement of the VLPFC in uncertain decision-making (Fellows and Farah, 2007), as well as in negative emotions (Vergallito *et al*., 2018) which could be linked to losing a round (i.e. the balloon exploding, hence, losing the points). In such context, the lack of sensitivity to emotions or uncertainty of the autoBART task, ill-adapted to capture the type of risk-taking that the VLPFC is in actively involved in, could explain the lack of modulatory effect under the effects of cathodal HD-tDCS, with the exception of participants with specific personality profiles.

Indeed, it is interesting to note that even if significant differences (with moderate effect sizes) vanished after Bonferroni’s correction, these pointed out in the same direction: the means of all dependent variables (but averaged adjusted pumps) in the sham condition reflected the expected trend from the personality profiling. *Impulsive-disinhibited* participants engaged in more risk-taking behaviour than *normative* participants, and the latter more than the *inhibited/emotional* profile. Taken together, such differences could relate to the larger tDCS modulatory effect that cathodal stimulation showed on personality Profile 1 compared to the two other profiles.

Further research will be needed to shed additional light on the indirect influence of personality differences on HD-tDCS impact. Nonetheless, in agreement with our main planned analysis, the post-hoc reanalysis of our dataset filtered by personality profiles also support the hypothesis of a differential role for DLPFC and VLPFC in risk decision-making. Moreover, such post-hoc analysis suggests that personality features could indirectly explain such dissociation. Specifically, our results provide a preliminary basis to argue that whereas the left VLPFC would be casually involved in increasing risk propensity in normative individuals, the left DLPFC might increase risk propensity in impulsive-disinhibited (profile 1) participants.

Regardless, the influence of personality profiles on the prefrontal contributions to risk decision-making and its modulation by tDCS should not come as a surprise. Previous studies have suggested that emotions, sensation-seeking impulses, psychopathy, and impulsiveness influence risk-taking behaviour (Hunt *et al*., 2005; Suhr and Tsanadis, 2007; Humphreys *et al*., 2013; Lauriola *et al*., 2013). In agreement, a post-hoc analysis of the sham tDCS condition performed in our study (hence in absence of any effective external manipulation of cortical excitability) points out that personality traits such as narcissism and psychopathy correlate with risk decision-making, a result that aligns with existing literature (Klayman *et al*., 1999; Campbell *et al*., 2004; Camchong *et al*., 2007). In fact, the ‘dark-triad’ is a good predictor of overconfidence (Wissing and Reinhard, 2017), which may ultimately explain why high scorers tend to take higher behavioural risks.

Our post-hoc analyses also suggest potential inferences on regions subtending personality differences, which could casually influence the impact of HD-tDCS on risk decision-making. Previous research has shown that DLPFC plays an important role not only in emotional regulation and risk behaviours (Kaiser *et al*., 2018) but also in aggression (Buckholtz and Meyer-Lindenberg, 2008). Hence the modulation of DLPFC activity with tDCS could show a particular impact in individuals with dysfunctional patterns of network interaction. To this regard, the “state-dependency” framework (Silvanto and Pascual-Leone, 2008) could provide a useful mechanistic framework to interpret our results. According to this theory, the ongoing activity and excitability estimates operating on a stimulated cortical area and its associated networks has the potential to influence the net sign and the magnitude of the stimulation effects by either TMS or tDCS. Seminal studies showed that when the motor cortical excitability was preconditioned using tDCS, this effect modulated the direction of the effects produced by later rTMS patterns (Siebner *et al*., 2004). Additional research with single pulses or repetitive TMS trains pointed out that in the visual cortex less active brain regions seem to be more susceptible to activating effects of single TMS pulses or rTMS trains (Silvanto *et al*., 2007). Furthermore, state dependency is a factor to take into consideration in order to plan, predict and eventually boost the impact of NIBS on brain areas (Silvanto and Pascual-Leone, 2008). Hence, we here argue that personality profiles might result in constitutional differences in ongoing prefrontal activity patterns and excitability estimates, explaining differences in the magnitude and direction of modulatory effects via NIBS. Further work would be necessary to characterize differences of activity state across personality profiles and confirm this hypothesis.

Alternatively, the observed influence of personality differences could be analysed by considering interindividual differences and deviations from the mean as ‘noise’ and a source of error. Unfortunately, very few studies to date have explored the role of interindividual differences to explain such error, which in the data is featured by unexplained variability (Zimerman and Hummel, 2010; Baeken *et al*., 2019; Qi *et al*., 2019). Although not always acknowledged in cognitive neuroscience, behavioural phenotypical variabilities in response to perturbations are subtended by inherent interindividual differences in age, plasticity (Zimerman and Hummel, 2010), brain structure (DeYoung *et al*., 2011; Li *et al*., 2015) and function (Corr, 2004), all of them likely to interfere with the neuromodulatory effects of non-invasive stimulation. Hence, the influence of personality profiles suggested by our study highlight the importance of personality neuroscience. Most importantly, it enacts an agenda to make use of tools used in cognitive and affective neuroscience to link brain processes to personality features (Markett *et al*., 2018) to ultimately identify the underlying neurobiological sources of personality (Allen and Deyoung, 2016), such as the case for personality dimensions like the Big Five Personality traits (Li *et al*., 2017).

### 4.3 Methodological considerations

Several methodological and design considerations need to be taken into account when interpreting the current results and envisioning future steps. First, the present study on the left prefrontal systems (DLPFC and VLPFC) was sensitive to gains and disregarded right homotopic prefrontal regions rather sensitive to losses (Ye, Chen, Huang, Wang and Luo, 2015) since the autoBART paradigm places an emphasis on the former rather than the latter. Hence, we cannot rule out if in retrospective, this decision might be at the base of the dissociation reported between the DLPFC and the VLPFC, justifying in our study the post-hoc exploration of personality profiles. Second, controversy exists with regards to the optimal tools to measure individual differences in personality. This is relevant given that tools are differentially sensitive to reward (e.g., Sensitivity to Punishment and Sensitivity to Reward Questionnaire SPSRQ (Torrubia, R., Avila, C., Molto, J., and Grande, 1995), the BIS/BAS scales (Carver and White, 1994) or the Scale for Measuring Reward Responsiveness (Van den Berg *et al*., 2010) and could have clustered our populations differently in personality types. Unfortunately, we lack the data to assess the robustness of our findings across different personality scales. Nonetheless, after considering different options, we adopted on risk-taking assessment by means of the HEXACO and the Dark Triad, a choice that optimized comparability with other psychological and behavioural research on risk-taking (Vries *et al*., 2009; Weller and Thulin, 2012; Pletzer *et al*., 2019; Maneiro *et al*., 2020). Third, our outcomes on the impact of personality traits on tDCS modulatory effects needs to be taken with caution, given the small and unequal samples of personality clusters considered in our post-hoc analyses (according to which *impulsive-disinhibited* profile counted with n=10 participants, *normative* profile n=18 and *inhibited/emotional* profile 3 included only n=6 participants). Most importantly, personality traits were not analysed pre-hoc and used to build equivalent experimental groups to be assigned to DLPFC or VLPFC tDCS, but rather applied as a *post-hoc* clustering criteria to assess its impact of regional prefrontal stimulation on risk taking behaviours.

Regardless, we hypothesize that some of our results could be replicated in prospective studies and larger and equal samples sharing personality profiles. This prediction is based on the fact that the observed power of 0.98 to correctly identify the number of profiles using 8 indicators, and a mean effect size between profiles of 0.53 identified with the SABIC, suggests a large effect in personality research (Gignac and Szodorai, 2016). Moreover, outcomes from the LPA and average effect size proved sound, replicable, and in survived power simulations, whereas Bayesian approaches were applied to overcome additional pitfalls.

Fourth and last, given prior research showing instances of more consistent outcomes for anodal than cathodal stimulation (Filmer *et al*., 2014; Lavidor, 2016; Thair *et al*., 2017), the choice of a cathodal tDCS -aimed at suppressing local excitability-could have also hindered our results. Additionally, the low magnitude of the induced electrical fields and the poor spatial resolution of tDCS and overlapping electric fields (Huang *et al*., 2019) could have also weakened functional dissociations between ventral and dorsal lateral prefrontal targets. Nonetheless, the low risk profile, ease of use, low cost, ability to produce reliable sham conditions and spread across extended subregions of prefrontal cortex, plus its convenience for therapeutic applications (e.g., such as in compulsive risk behaviours) made tDCS the most suited choice of NIBS technology for our study. Provided that specific MNI xyz MRI/fMRI coordinates for DLPFC and VLPFC targets involved in the autoBART become available, a replication of our experiment with a more powerful and higher spatial resolution NIBS approach such as TMS is advised.

## 5. CONCLUSIONS

We carried out a study to assess the dorsal and ventral prefrontal contributions to risk-taking behaviours as assessed by the autoBART task. We found that cathodal HD-tDCS delivered to the prefrontal left DLPFC but not the VLPFC modified risk-taking behaviour. Importantly, a re-analysis of the same data revealed that such prefrontal subregion-dependent effects are influenced by the personality profiles participants were clustered into according to the HEXACO and Dark Triad dirty dozen scores. More specifically, modulation of risk propensity with left DLPFC HD-tDCS affected impulsive-disinhibited individuals, whereas left VLPFC stimulation influenced risk propensity in normative participants. On such basis, we conclude that a systematic profiling of personality-based differences may offer insights into the anatomical basis of normal and abnormal personality traits and might help identify and select pre-hoc which individuals will be most suited to respond to NIBS interventions.

## Abbreviations

(AVP): adjusted average of pumps
(autoBART): auto balloon analogue risk task
(DLPFC): dorsolateral prefrontal cortex
(HD-tDCS): high-definition transcranial direct current stimulation
(TE): total earnings
(tDCS): transcranial direct current stimulation
(TMS): transcranial magnetic stimulation
(VLPFC): ventrolateral prefrontal cortex
(WP): wanted pumps

## Footnotes

1 Mean Cohen’s d between classes (expressed as multivariate Mahalanobis’ distance) was 0.53. Considering this effect size, our sample size (n=34), and the number of indicators used (k=8) to classify participants, we run a number of simulations to estimate our observed power. Results indicated that SABIC yielded correct classifications in 98% of replications (after 1,000 replications). In small-size samples, SABIC clearly outperforms BIC (in these simulations, BIC yielded correct classifications only in 17% of replications (Gallardo-Pujol, in preparation). Scripts and data of the simulations are available here https://osf.io/m79pg/?view_only=887f5b53d3b14319898afac7b7391885.

