## Supplementary material for "Modulatory effect of non-invasive prefrontal brain stimulation on risk-taking behaviours in humans: preliminary insight on the influence of personality traits"

**Table S.1 Descriptive analysis of personality data**

| Personality measures | N | Min | Max | Mean | Std. Deviation |
|---|---|---|---|---|---|
| Narcissism | 34 | 4.00 | 20.00 | 9.73 | 3.74 |
| Psychopathy | 34 | 4.00 | 15.00 | 8.09 | 2.97 |
| Machiavellianism | 34 | 4.00 | 16.00 | 10.09 | 3.83 |
| Openness to Experience | 34 | 27.00 | 47.00 | 37.15 | 5.18 |
| Conscientiousness | 34 | 27.00 | 49.00 | 37.35 | 5.49 |
| Agreeableness | 34 | 17.00 | 49.00 | 31.47 | 7.59 |
| Extraversion | 34 | 15.00 | 46.00 | 32.70 | 7.43 |
| Emotionality | 34 | 19.00 | 46.00 | 32.03 | 6.73 |
| Honesty/Humility | 34 | 20.00 | 46.00 | 33.76 | 7.27 |

**Table S.2 Correlations between personality and dependent variables. Correlations between personality and dependent variables.** HEXACO and dark-triad correlations with the BART dependent variables.

1) BART and personality dimensions

1a) Average adjusted pumps and HEXACO

Descriptive statistics (N=34)

| Variables | 1 | 2 | 3 | 4 | 5 | 6 | 7 | 8 | 9 |
|---|---|---|---|---|---|---|---|---|---|
| 1.Average adjusted pumps DLPFC | - | .85** | .90** | -.16 | -.08 | .33 | -.03 | -.40* | -.12 |
| 2. Average adjusted pumps VLPFC | .85** | - | .88** | -.31 | -.15 | .23 | .11 | - .43** | -.23 |
| 3. Average adjusted pumps Sham | .90** | .88** | - | -.13 | .01 | .23 | .05 | -.41* | -.06 |
| 4.Openness to Experience | -.16 | -.31 | -.13 | - | .48** | -.07 | -.19 | -.10 | .30 |

| | | | | | | | | | |
|---|---|---|---|---|---|---|---|---|---|
| 5.Conscientiousness | -.08 | -.15 | .01 | .48** | - | .04 | .09 | -.25 | .35* |
| 6.Agreeableness | .33 | .23 | .23 | -.07 | .04 | - | .11 | -.24 | .19 |
| 7.Extraversion | -.03 | .11 | .05 | -.19 | .09 | .11 | - | -.36* | -.21 |
| 8.Emotionality | -.40* | -.43** | -.41* | -.10 | -.25 | -.24 | -.36* | - | .08 |
| 9.Honesty/Humility | -.12 | -.23 | -.06 | .30 | .35* | .19 | -.21 | .08 | - |

*Note*. *p < .05. **p < .01.

Continued

| Variables | 1 | 2 | 3 | 4 | 5 | 6 | 7 | 8 | 9 |
|---|---|---|---|---|---|---|---|---|---|
| M | 50.69 | 51.37 | 54.04 | 37.15 | 37.35 | 31.47 | 32.70 | 32.03 | 33.76 |
| SD | 13.26 | 16.43 | 16.39 | 5.18 | 5.49 | 7.59 | 7.43 | 6.73 | 7.27 |

1b) Average adjusted pumps and dark triad

Descriptive statistics (N=34)

| Variables | 1 | 2 | 3 | 4 | 5 | 6 |
|---|---|---|---|---|---|---|
| 1. Average adjusted pumps DLPFC | - | .85** | .90** | .13 | .47** | -.10 |
| 2. Average adjusted pumps VLPFC | .85** | - | .88** | .14 | .44** | -.06 |
| 3. Average adjusted pumps Sham | .90** | .88** | - | .02 | .44** | -.07 |
| 4.Machiavellianism | .13 | .14 | .02 | - | .49** | .49** |
| 5.Psychopathy | .47** | .44** | .44** | .49** | - | .14 |
| 6.Narcissism | -.10 | -.06 | -.07 | .49** | .14 | - |

*Note*. *p < .05. **p < .01.

Continued

| Variables | 1 | 2 | 3 | 4 | 5 | 6 |
|---|---|---|---|---|---|---|
| M | 50.69 | 51.37 | 54.04 | 10.09 | 8.09 | 9.73 |
| SD | 13.26 | 16.43 | 16.39 | 3.83 | 2.97 | 3.74 |

1c) Total earnings and HEXACO

Descriptive statistics (N=34)

| Variables | 1 | 2 | 3 | 4 | 5 | 6 | 7 | 8 | 9 |
|---|---|---|---|---|---|---|---|---|---|
| 1.Total Earnings DLPFC | - | .25 | -.21 | -.13 | -.25 | .22 | .16 | -.22 | -.35* |

| | | | | | | | | | |
|---|---|---|---|---|---|---|---|---|---|
| 2. Total Earnings VLPFC | .25 | - | .32 | -.37* | -.50** | -.01 | -.26 | .05 | -.38* |
| 3. Total Earnings Sham | -.21 | .32 | - | -.04 | .14 | -.26 | -.12 | .08 | -.16 |
| 4.Openness to Experience | -.13 | -.37* | -.04 | - | .48** | -.07 | -.19 | -.10 | .30 |
| 5.Conscientiousness | -.25 | -.50** | .14 | .48** | - | .04 | .09 | -.25 | .35* |
| 6.Agreeableness | .22 | -.01 | -.26 | -.07 | .04 | - | .11 | -.24 | .19 |
| 7.Extraversion | .16 | -.26 | -.12 | -.19 | .09 | .11 | - | -.36* | -.21 |
| 8.Emotionality | -.22 | .05 | .08 | -.10 | -.25 | -.24 | -.36* | - | .08 |
| 9.Honesty/Humility | -.35* | -.38* | -.16 | .30 | .35* | .19 | -.21 | .08 | - |

*Note*. *p < .05. **p < .01.

Continued

| Variables | 1 | 2 | 3 | 4 | 5 | 6 | 7 | 8 | 9 |
|---|---|---|---|---|---|---|---|---|---|
| M | 792.47 | 796.24 | 826.38 | 37.15 | 37.35 | 31.47 | 32.70 | 32.03 | 33.76 |
| SD | 190.30 | 176.80 | 171.01 | 5.18 | 5.49 | 7.59 | 7.43 | 6.73 | 7.27 |

1d) Total earnings and dark triad

Descriptive statistics (N=34)

| Variables | 1 | 2 | 3 | 4 | 5 | 6 |
|---|---|---|---|---|---|---|
| 1. Total Earnings DLPFC | - | .25 | -.21 | .32 | .39* | -.01 |
| 2. Total Earnings VLPFC | .25 | - | .32 | .35* | .32 | .03 |
| 3. Total Earnings Sham | -.21 | .32 | - | .01 | .19 | .18 |
| 4.Machiavellianism | .32 | .35* | .01 | - | .49** | .49** |
| 5.Psychopathy | .39* | .32 | .19 | .49** | - | .14 |
| 6.Narcissism | -.01 | .03 | .18 | .49** | .14 | - |

*Note*. *p < .05. **p < .01.

Continued

| Variables | 1 | 2 | 3 | 4 | 5 | 6 |
|---|---|---|---|---|---|---|
| M | 792.47 | 796.24 | 826.38 | 10.09 | 8.09 | 9.73 |
| SD | 190.30 | 176.80 | 171.01 | 3.83 | 2.97 | 3.74 |

1e) Total wanted pumps and HEXACO

Descriptive statistics (N=34)

| Variables | 1 | 2 | 3 | 4 | 5 | 6 | 7 | 8 | 9 |
|---|---|---|---|---|---|---|---|---|---|
| 1.Total Wanted pumps DLPFC | - | .84** | .91** | -.04 | .00 | .30 | -.03 | -.36* | -.03 |
| 2. Total Wanted pumps VLPFC | .84** | - | .83** | -.28 | -.07 | .21 | .13 | -.42* | -.14 |
| 3. Total Wanted pumps Sham | .91** | .83** | - | -.01 | .02 | .24 | .12 | -.38* | -.01 |
| 4.Openness to Experience | -.04 | -.28 | -.01 | - | .48** | -.07 | -.19 | -.10 | .30 |
| 5.Conscientiousness | .00 | -.07 | .02 | .48** | - | .04 | .09 | -.25 | .35* |
| 6.Agreeableness | .30 | .21 | .24 | -.07 | .04 | - | .11 | -.24 | .19 |
| 7.Extraversion | -.03 | .13 | .12 | -.19 | .09 | .11 | - | -.36* | -.21 |
| 8.Emotionality | -.36* | -.42* | -.38* | -.10 | -.25 | -.24 | -.36* | - | .08 |
| 9.Honesty/Humility | -.03 | -.14 | -.01 | .30 | .35* | .19 | -.21 | .08 | - |

*Note*. *p < .05. **p < .01.

Continued

| Variables | 1 | 2 | 3 | 4 | 5 | 6 | 7 | 8 | 9 |
|---|---|---|---|---|---|---|---|---|---|
| M | 1733.94 | 1705.18 | 1820.65 | 37.15 | 37.35 | 31.47 | 32.70 | 32.03 | 33.76 |
| SD | 412.47 | 474.27 | 571.66 | 5.18 | 5.49 | 7.59 | 7.43 | 6.73 | 7.27 |

1f) Total wanted pumps and dark triad

Descriptive statistics (N=34)

| Variables | 1 | 2 | 3 | 4 | 5 | 6 |
|---|---|---|---|---|---|---|
| 1. Total Wanted pumps DLPFC | - | .84** | .91** | .01 | .39* | -.09 |
| 2. Total Wanted pumps VLPFC | .84** | - | .83** | .10 | .38* | -.05 |
| 3. Total Wanted pumps Sham | .91** | .83** | - | .01 | .37* | -.06 |
| 4.Machiavellianism | .01 | .10 | .01 | - | .49** | .49** |
| 5.Psychopathy | .39* | .38* | .37* | .49** | - | .14 |
| 6.Narcissism | -.09 | -.05 | -.06 | .49** | .14 | - |

*Note*. *p < .05. **p < .01.

Continued

| Variables | 1 | 2 | 3 | 4 | 5 | 6 |
|---|---|---|---|---|---|---|
| M | 1733.94 | 1705.18 | 1820.65 | 10.09 | 8.09 | 9.73 |
| SD | 412.47 | 474.27 | 571.66 | 3.83 | 2.97 | 3.74 |

**Table S.3.** Pairwise comparison of cathodal DLPFC vs sham, cathodal VLPFC vs sham and cathodal VLPFC vs cathodal DLPFC under different personality groups.

| | | | | t | Mean Difference | Std. Error | Sig.[a] | 95% CI |
|---|---|---|---|---|---|---|---|---|
| **Average adjusted Pumps** | **Profile 1 (n=10)** | **DLPFC** | **VLPFC** | -.037 | -4.628 | 4.604 | .970 | -16.352-7.096 |
| | | | **Sham** | -2.05 | -9.305 | 3.655 | .050 | -18.613-.004 |
| | | **VLPFC** | **DLPFC** | .037 | 4.628 | 4.604 | .970 | -7.096-16.352 |
| | | | **Sham** | -.155 | -4.676 | 6.361 | .878 | -15.783-6.430 |
| | | **Sham** | **DLPFC** | 2.05 | 9.305 | 3.655 | .050 | -.004-18.613 |
| | | | **VLPFC** | .155 | 4.676 | 4.361 | .878 | 6.430-15.783 |
| | **Profile 2 (n=18)** | **DLPFC** | **VLPFC** | - | .181 | 2.072 | 1.000 | -5.095-5.456 |
| | | | **Sham** | -1.92 | -4.003 | 1.645 | .065 | -8.192-.185 |
| | | **VLPFC** | **DLPFC** | - | .181 | 2.072 | 1.000 | -5.456-5.095 |
| | | | **Sham** | -1.57 | -4.184 | 1.963 | .126 | -9.182-.814 |
| | | **Sham** | **DLPFC** | 1.92 | 4.003 | 1.645 | .065 | -.185-8.192 |
| | | | **VLPFC** | 1.57 | 4.184 | 1.963 | .126 | -.814-9.182 |
| | **Profile 3 (n=6)** | **DLPFC** | **VLPFC** | - | .151 | 3.566 | 1.000 | -8.930-9.232 |
| | | | **Sham** | - | -.592 | 2.831 | 1.000 | -7.802-6.618 |
| | | **VLPFC** | **DLPFC** | - | -.151 | 3.566 | 1.000 | -9.232-8.930 |
| | | | **Sham** | - | -.743 | 3.378 | 1.000 | -9.346-7.860 |

| | | | | | | | |
|---|---|---|---|---|---|---|---|
| | | **Sham** | **DLPFC** | - | .592 | 2.831 | 1.000 | -6.618-7.802 |
| | | | **VLPFC** | - | .743 | 3.378 | 1.000 | -7.860-9.346 |
| **Wanted Pumps** | **Profile 1 (n=10)** | **DLPFC** | **VLPFC** | -.051 | -176.056 | 135.228 | .611 | -520.409-168.298 |
| | | | **Sham** | -1.99 | -322.278 | 129.175 | .056 | -651.217-6.661 |
| | | **VLPFC** | **DLPFC** | .051 | 176.056 | 135.228 | .611 | -168.298-520.409 |
| | | | **Sham** | - | -146.222 | 179.651 | 1.000 | -603.697-311.253 |
| | | **Sham** | **DLPFC** | 1.99 | 322.278 | 129.175 | .056 | -6.661-651.217 |
| | | | **VLPFC** | - | 146.222 | 179.651 | 1.000 | -311.253-603.697 |
| | **Profile 2 (n=18)** | **DLPFC** | **VLPFC** | - | 49.100 | 60.853 | 1.000 | -105.859-204.059 |
| | | | **Sham** | -.62 | -80.087 | 58.129 | .538 | -228.110-67.935 |
| | | **VLPFC** | **DLPFC** | - | -49.100 | 60.853 | 1.000 | -204.059-105.859 |
| | | | **Sham** | -.092 | -129.187 | 80.843 | .364 | -335.051-76.676 |
| | | **Sham** | **DLPFC** | .62 | 80.087 | 58.129 | .538 | -67.935-228.110 |
| | | | **VLPFC** | .092 | 129.187 | 80.843 | .364 | -76.676-335.051 |
| | **Profile 3 (n=6)** | **DLPFC** | **VLPFC** | .060 | 106.667 | 104.747 | .952 | -160.068-373.402 |
| | | | **Sham** | - | 58.833 | 100.058 | 1.000 | -195.962-313.628 |
| | | **VLPFC** | **DLPFC** | -0.06 | -106.667 | 104.747 | .952 | -373.402-160.068 |
| | | | **Sham** | - | -47.833 | 139.157 | 1.000 | -402.192-306.525 |
| | | **Sham** | **DLPFC** | - | -58.833 | 100.058 | 1.000 | -313.628-195.962 |
| | | | **VLPFC** | - | 47.833 | 139.157 | 1.000 | -306.525-402.192 |
| | | | **VLPFC** | -.27 | -127.833 | 111.820 | .788 | -412.578-156.911 |

| | | | | | | | | |
|---|---|---|---|---|---|---|---|---|
| | | **DLPFC** | **Sham** | - | -88.611 | 147.528 | 1.000 | -464.287-287.065 |
| | **Profile 1 (n=10)** | **VLPFC** | **DLPFC** | .27 | 127.833 | 111.820 | .788 | -156.911-412.578 |
| | | | **Sham** | - | 39.222 | 103.050 | 1.000 | -223.190-301.635 |
| | | **Sham** | **DLPFC** | - | 88.611 | 147.528 | 1.000 | -287.065-464.287 |
| | | | **VLPFC** | - | -39.222 | 103.050 | 1.000 | -301.635-223.190 |
| **Total Earnings BART** | **Profile 2 (n=18)** | **DLPFC** | **VLPFC** | - | 14.012 | 50.319 | 1.000 | -114.123-142.148 |
| | | | **Sham** | -.48 | -84.987 | 66.388 | .633 | -254.042-84.067 |
| | | **VLPFC** | **DLPFC** | - | -14.012 | 50.319 | 1.000 | -142.148-114.123 |
| | | | **Sham** | -1.58 | -99.000 | 46.372 | .125 | -217.086-19.086 |
| | | **Sham** | **DLPFC** | .48 | 84.987 | 66.388 | .633 | -84.067-254.042 |
| | | | **VLPFC** | 1.58 | 99.000 | 46.372 | .125 | -19.086-217.086 |
| | **Profile 3 (n=6)** | **DLPFC** | **VLPFC** | -1.05 | -147.167 | 86.615 | .301 | -367.729-73.396 |
| | | | **Sham** | - | -73.000 | 114.275 | 1.000 | -363.997-217.997 |
| | | **VLPFC** | **DLPFC** | 1.05 | 147.167 | 86.615 | .301 | -73.396-367.729 |
| | | | **Sham** | - | 74.167 | 79.822 | 1.000 | -129.097-277.430 |
| | | **Sham** | **DLPFC** | - | 73.000 | 114.275 | 1.000 | -217.997-363.997 |
| | | | **VLPFC** | - | -74.167 | 79.822 | 1.000 | -277.430-129.097 |

**Note.** * First column, dependent variables; second, independent variables (personality profiles). a. Adjustment for multiple comparison: Bonferroni. DLPFC: Dorsolateral Prefrontal Cortex, VLPFC: Ventrolateral Prefrontal Cortex. The shaded rows are where statistically significant differences have been found.
